# Sensitivity to naturalistic texture relies primarily on high spatial frequencies

**DOI:** 10.1101/2022.08.23.504875

**Authors:** Justin D. Lieber, Gerick M. Lee, Najib J. Majaj, J. Anthony Movshon

## Abstract

Natural images contain information at multiple spatial scales. Although we understand how early visual mechanisms split multi-scale images into distinct spatial frequency channels, we do not know how the outputs of these channels are processed further by mid-level visual mechanisms. We have recently developed a naturalness discrimination task that uses synthesized, multi-scale textures to isolate these mid-level mechanisms (Freeman et. al. 2013). Here, we use three experimental manipulations (image blur, image rescaling, and eccentric viewing) to show that naturalness sensitivity is strongly dependent on image features at high *object* spatial frequencies (measured in cycles/image). As a result, sensitivity depends on a *texture acuity limit*, a property of the visual system that sets the highest *retinal* spatial frequency (measured in cycles/degree) that can be used to solve the task. A model observer analysis shows that high object spatial frequencies carry more task-relevant information than low object spatial frequencies. Comparing the outcome of this analysis with human performance reveals that human observers’ efficiency is similar for all object spatial frequencies. We conclude that the mid-level mechanisms that underlie naturalness sensitivity effectively extract information from all image features below the texture acuity limit, regardless of their retinal and object spatial frequency.

## Introduction

Decades of research have produced strong, quantitative models of the early visual mechanisms that extract information from visual scenes. Studies in which observers report the presence of near-threshold stimuli indicate that the initial stages of visual processing split incoming signals into distinct channels selective for different spatial frequencies (Graham, 1989). These spatial frequency channels bear a resemblance to neural responses in primary visual cortex (V1), where individual neurons also respond selectively to specific spatial frequencies (De Valois et al., 1982).

Though successful for near-threshold experiments, early channel models have proven insufficient to explain the mid-level mechanisms that govern the perception of more complex, suprathreshold visual scenes. Because early mechanisms are primarily sensitive to spatial frequency amplitude, one way to neutralize their contribution is to generate sets of images that are matched in frequency amplitude (*“spectrally-matched”*), yet are nonetheless perceptually distinct (Chubb & Sperling, 1988). Using this technique, many groups have used spectrally-matched textures to study the detection of local modulations in features like contrast, orientation, or spatial frequency (reviewed in Graham, 2011). Controlling for early channel mechanisms means that these tasks provide insight into the “mid-level” visual mechanisms that lie beyond early visual processing.

A limit of these studies is that they typically used stimuli that 1) only modulated one specific feature (contrast, orientation, etc.) at a time and 2) were limited to a narrow band of spatial frequencies. As such, we still have an incomplete knowledge of how observers combine information from overlapping modulations in the same image (but see Landy & Kojima (2001), Saarela & Landy (2012)) and from information distributed across a broad range of spatial frequencies. These are both characteristic features of natural scenes.

An alternative approach to studying visual form has focused on the ability of observers to identify everyday objects and scenes (Morrison & Schyns, 2001; Sowden & Schyns, 2006). However, this approach comes with challenges. First, task-relevant information may be unevenly distributed across different spatial frequencies in the image. For example, discriminating two very different objects (i.e. dogs vs. cats) relies on lower spatial frequencies than discriminating two very similar objects (i.e. two breeds of dogs) (Archambault et al., n.d.). Experimenters have quantified this distribution of information by studying how task performance changes for images that have been low-pass or bandpass filtered to remove certain frequencies (Braje et al., 1995; Gold et al., 1999; Parish & Sperling, 1991).

A second challenge is distinguishing whether task performance is set by factors intrinsic to the images (measured in image-centric coordinates) or factors intrinsic to the visual system (measured in retina-centric coordinates). Factors intrinsic to the images are best described by *object spatial frequencies* that are measured as the number of *cycles per object*. This is categorically different than factors of the visual system, which are best described by *retinal spatial frequencies* that are measured in *cycles per degree*. For example, one particularly well studied limit of the visual system is the contrast sensitivity function, which describes how the visibility of image features changes with retinal spatial frequency and location in visual field (Anderson et al., 1991; Robson & Graham, 1981). How an image’s object spatial frequencies correspond to an observer’s retinal spatial frequencies is determined by the distance at which the observer views the image. At close distances, images take up more of the visual field and object spatial frequencies correspond to low retinal spatial frequencies. At far distances, images become increasingly small and object spatial frequencies correspond to high retinal spatial frequencies. Manipulating viewing distance can dissociate whether task performance is set by factors intrinsic to the image (object spatial frequencies) or intrinsic to the visual system (retinal spatial frequencies) (Parish & Sperling, 1991).

One final challenge of using broadband images is evaluating the complexity of the image features that drive task performance. We can broadly group these features into three “levels” of complexity. First, a task could be solved solely with simple spectral cues, corresponding to the outputs of the early channel mechanisms. Second, a task might depend on features that are a simple, direct transformation of the outputs of the early channel stage, corresponding to “mid-level” mechanisms. Finally, a task might require complex features that can only be extracted after multiple levels of hierarchical transformation, corresponding to “late” mechanisms. The ability of human observers to readily distinguish natural images from spectrally-matched counterparts (Oppenheim & Lim, 1981; Piotrowski & Campbell, 1982; Thomson et al., 2000) demonstrates that natural scenes engage more than just early mechanisms. However, distinguishing whether mid-level or late mechanisms are used to solve a task requires a model of the specific features that mid-level mechanisms might extract.

We have recently developed a psychophysical framework that isolates the contribution of mid-level visual mechanisms by using synthesized texture images that capture many properties of natural texture (Freeman et al., 2013). In this context, we define a “texture” as a self-similar image defined by elements that repeat, possibly with variation (Graham & Landy, 2002; Landy, 2013). Many aspects of a texture’s self-similarity can be captured with statistical measures based on the proposed structure of mid-level vision (Heeger & Bergen, 1995; Portilla & Simoncelli, 2000). We used the Portilla & Simoncelli texture synthesis procedure to generate “naturalistic” texture images based on the statistics of photographs of natural texture. Naturalistic textures and spectrally-matched noise images drive similar levels of activity in V1, but downstream visual areas (like extrastriate areas V2 and V4) exhibit stronger responses to naturalistic texture images than to spectrally-matched noise (Freeman et al., 2013; Movshon & Simoncelli, 2014; Okazawa et al., 2015, 2016). The strength of this naturalness modulation varies between images with different texture statistics, and this variation predicts differences in perceptual sensitivity to naturalistic structure in those same images (Freeman et al., 2013).

Experiments using synthesized naturalistic textures combine the strengths of using parametrically defined texture stimuli with the strengths of discriminating natural scenes. Like other parametrically defined texture stimuli, naturalistic textures are defined by a discrete, predefined set of mid-level image features. Like natural objects, naturalistic textures contain a many overlapping mid-level image features distributed over a range of spatial frequencies. Naturalistic textures allow for the controlled investigation of mid-level visual processing in complex, multi-scale images.

In this study, we report measurements on how naturalness sensitivity changes in response to three experimental manipulations: image blur, image rescaling (equivalent to changes in viewing distance), and presentation of images in the peripheral visual field. Measurements from all three manipulations demonstrate that naturalness sensitivity is set primarily by two factors. First, observers are bound by a *texture acuity limit* (measured in cycles/degree) such that image features present in *retinal* spatial frequencies above this limit cannot be used to solve the naturalness task. Second, each family of texture images is associated with its own *cumulative sensitivity function* that describes how observers extract naturalness information as increasingly high *object* spatial frequencies (measured in cycles per image) are added to the image. For all texture families, cumulative sensitivity functions rise with object spatial frequency, demonstrating that high object frequencies are necessary to achieve high naturalness sensitivity. Finally, we find that a model observer analysis, based purely on the intrinsic informativeness of the image sets, closely tracks the cumulative sensitivity functions derived from measurement made on human observers.

## Experimental methods

### Image generation

We used the Portilla & Simoncelli analysis/synthesis procedure (Portilla & Simoncelli, 2000) to generate images of naturalistic texture. The procedure first measures the statistical properties of texture images, then synthesizes new images that match those statistical properties. For source images, we chose a set of 5 representative texture families from a larger set of 479 texture families used in a previous study (Freeman et al., 2013). Each texture family consisted of multiple synthesized images based on the statistics of a single black and white photograph of a visual texture. We chose families to maximally vary along the perceptual dimensions of coarse versus fine (the presence of low vs. high spatial frequencies), directional versus non-directional (whether orientations are highly concentrated or more evenly distributed), and regular versus irregular (whether patterns in the texture repeat at consistent or inconsistent intervals) (Rao & Lohse, 1996), quantified using methods described in Kim et al. (2019, 2022).

We then used the Portilla/Simoncelli analysis/synthesis procedure to measure the statistical properties of these textures images by first computing the output of a set of V1-like filters that can differ in orientation, spatial frequency, and location (Figure 1A-B). Two sets of filters capture linear (simple-cell like) and rectified (energy, complex-cell like) outputs. These ensembles of filters are then multiplied and averaged to yield cross-orientation, cross-scale, and cross-location covariance statistics (Figure 1C). The procedure also measures and matches the average energy, skew, and kurtosis of each filter ensemble.

**Figure 1.**
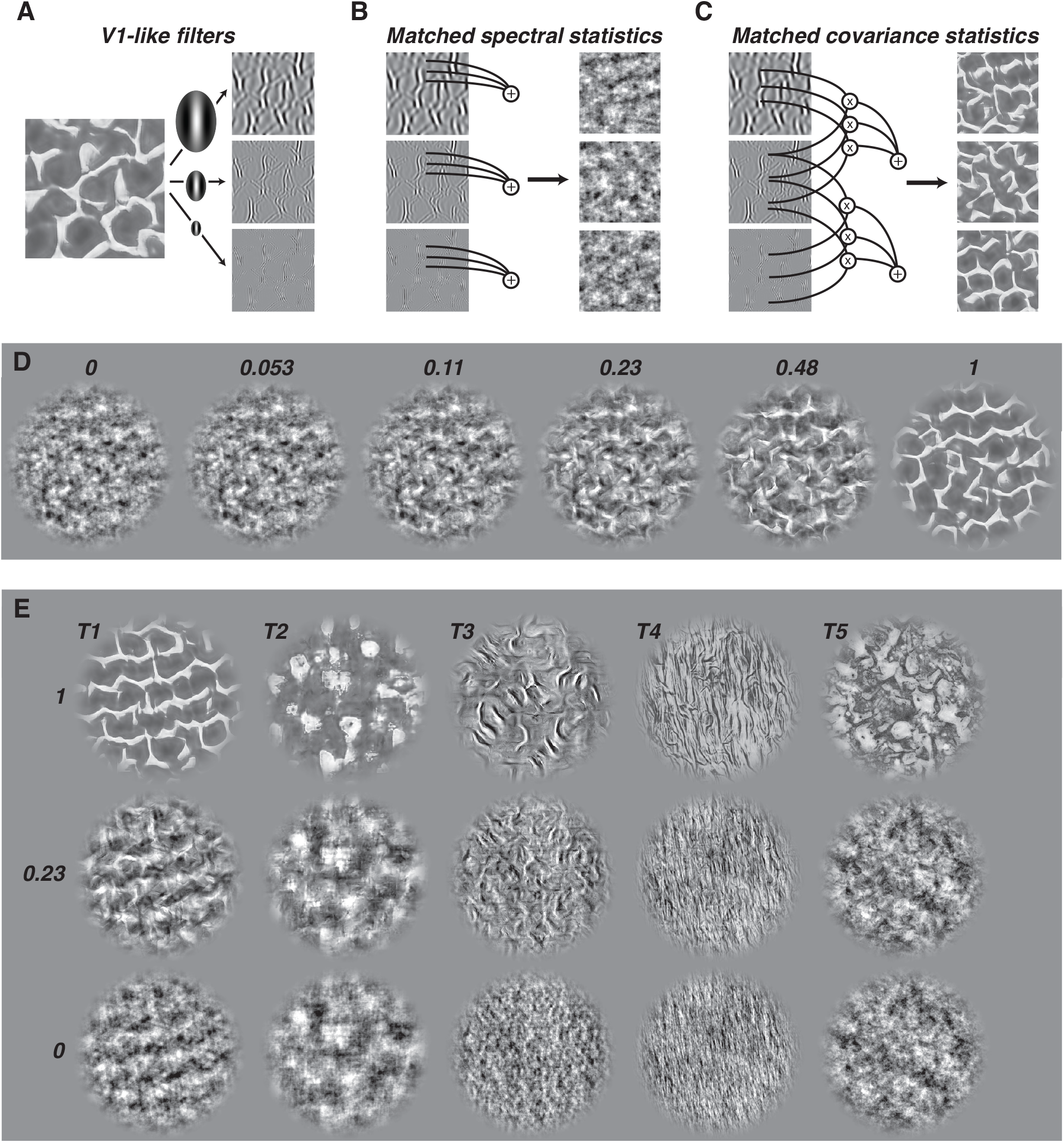
Synthesis of naturalistic textures. A) Texture images are filtered into subbands centered at different orientations and spatial frequencies. B) Measuring the average activation within each subband (left) creates a set of statistics that summarize the amplitude spectrum of the original image. Synthesizing new images that match these statistics creates “noise” textures that are spectrally-matched to the original sample. Three different random initializations of the synthesis creates three distinct spectrally-matched images. C) Naturalistic texture images match both the spectral statistics of the original image as well as products of filter responses across different spatial frequency and orientation bands (left). The resulting texture images (right) contain many of the naturalistic features found in natural texture images. D) Example texture images of intermediate naturalness. The textures shown have naturalness levels of 0, 0.053, 0.11, 0.23, 0.48, and 1, from left to right. E) Examples from the five texture families used in this study, labeled T1 to T5. The textures shown have naturalness levels of 1, 0.23, and 0, from top to bottom.

To synthesize new images, we first created “noisy” seed images by phase-randomizing the Fourier spectrum of the original texture images. This creates images that match the spectral content of the original texture images but lack higher-order correlations and naturalistic structure. We then adjusted the statistics of these noise images (using gradient descent) to match the model parameters of the original texture images. For each family, running this process on 15 distinct phase-randomized images yielded 15 distinct naturalistic texture images.

To generate images of intermediate naturalness (Figure 1D) that smoothly transition between fully naturalistic texture images and spectrally-matched noise we linearly interpolated model parameters between naturalistic statistics and noise statistics. For model parameters *p_orig_* for the original images and *p_noise_* for noise images, we computed:

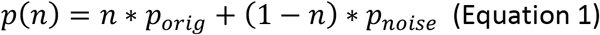

where *n* is the “naturalness” level of the statistics. Images were then synthesized to match these statistics, as above, initialized using the same base set of 15 phase-randomized noise images. We chose naturalness levels to be 17 logarithmically distributed values between 0.02 and 1 (0.353 octaves between points), plus one additional level at 0. The resulting images for the 5 texture families (Figure 1E) were used in all naturalness experiments.

Of particular importance to this set of experiments are the limits of the spatial frequency bands used in the synthesis model. All bands span a range of object spatial frequencies 1 octave wide. The images used in this study were 320 pixels wide, resulting in a Nyquist frequency *f_nyq_* = 160 cycles/image. The highest frequency band in the synthesis model spans a range of object spatial frequencies from 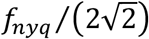 to 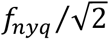 (Figure 2). Thus, the upper bound of the highest object spatial frequency band in our study, which we refer to as the *synthesis limit* (*f_synth_*), is 113 cycles/image.

**Figure 2.**
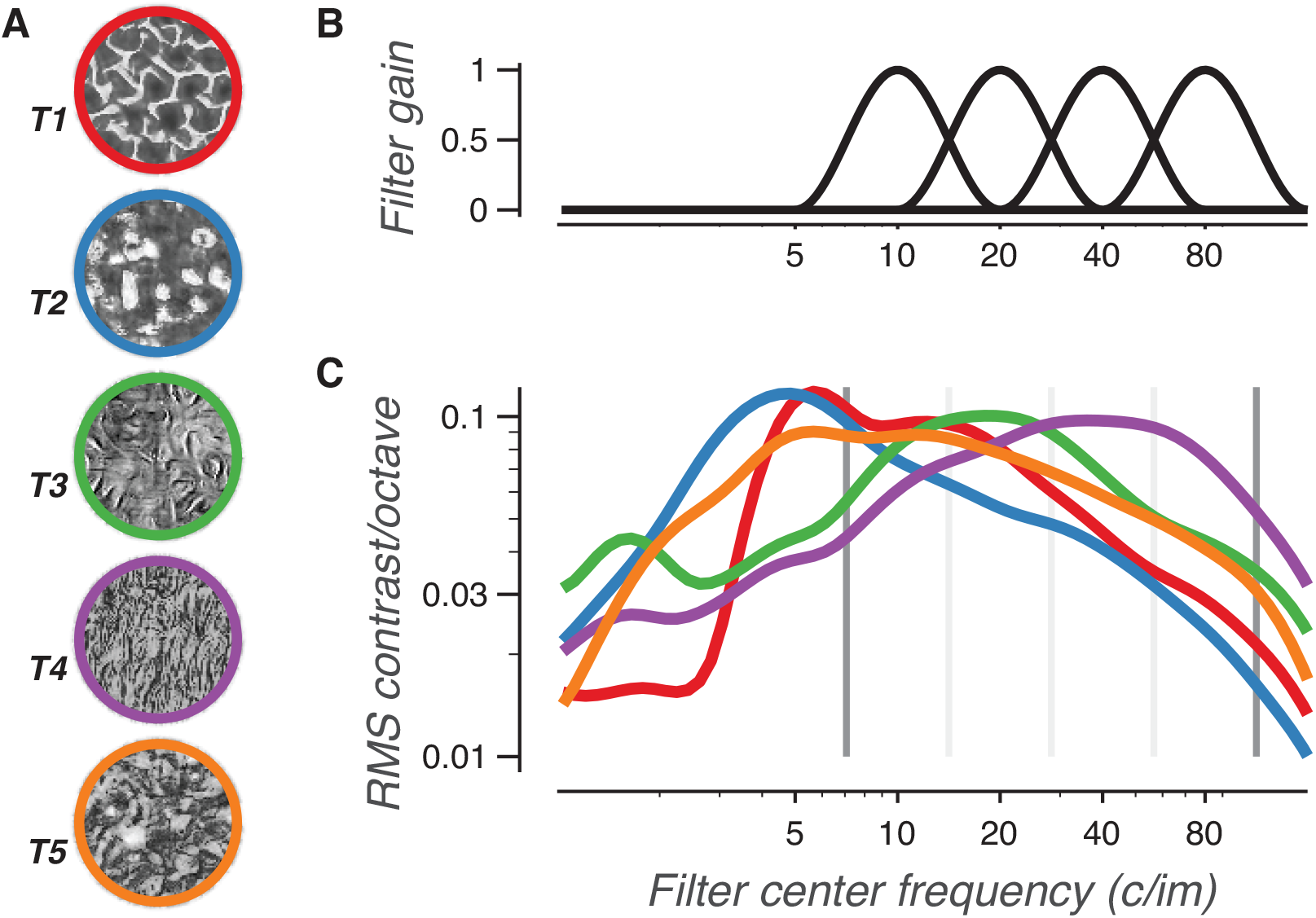
Distribution of contrast over object spatial frequency bands for different texture families. A) Example images of the five texture families. B) The four curves plot the spatial frequency filters used to create the bands of the texture synthesis model. C) Curves plot the average contrast of bandpass filtered images from each texture family. All naturalistic texture and spectrally-matched noise images in each family were filtered by bandpass filters with a width of one octave (same shape as those used in the texture synthesis model) centered at different spatial frequencies. RMS contrast was averaged over the resulting images. Note that because the filters were of even width in log frequency space (and of increasing width in linear frequency space), an image set with a 1/f amplitude spectra would plot as a flat, horizontal line. Grey lines denote the edges of the four frequency bands. Dark grey lines represent the lowest and highest spatial frequencies used to measure texture statistics for the synthesis model (3.5 and 113 cycles/image), and thus the range of object spatial frequencies that can be used to solve the naturalness sensitivity task.

### Psychophysical methods

3 male observers (age 23-36) with normal or corrected-to-normal vision participated in the experiments. 2 observers were authors. The third observer was naïve to the purpose of the study. Psychophysical results from the naïve observer (data/fits in purple throughout the paper) were indistinguishable from those of the two authors. Protocols for the experimental procedures were approved by the Institutional Review Board of New York University.

Image pixel luminance was rescaled to achieve an RMS contrast of 0.2, then vignetted in circular patches of 320 pixel diameter (flat top, raised cosine edges with width = 20 pixels). Images were presented on a 40 by 30 cm flat screen CRT monitor at a distance of 142 cm, unless otherwise noted. The resolution of the screen was 1280 × 960 pixels, which at 142 cm corresponds to a resolution of 80 pixels/deg. Thus, at this distance, images spanned 4 degrees of visual angle. Unless otherwise noted, observers were always instructed to look directly at the images.

Observers performed a three-interval, two-alternative forced choice, match-to-sample task that measured their ability to discriminate images of naturalistic texture from images of spectrally-matched noise (Figure 3). On every trial, three images were flashed in quick succession (200 ms presentation time, 300 ms inter-presentation intervals). The first and last images presented were constrained to be, in either order, one sample of spectrally-matched noise and one sample of naturalistic texture. The middle image was a distinct image that matched the naturalness level of either the first or last image. Observers had to report which pair of images belonged to a matching category: either the first two images or last two images. The primary benefit of this task structure is that it imposes no assumptions on observers as to what image features should be used to solve the task (Hillis et al., 2002). Observers received feedback after every trial.

**Figure 3.**
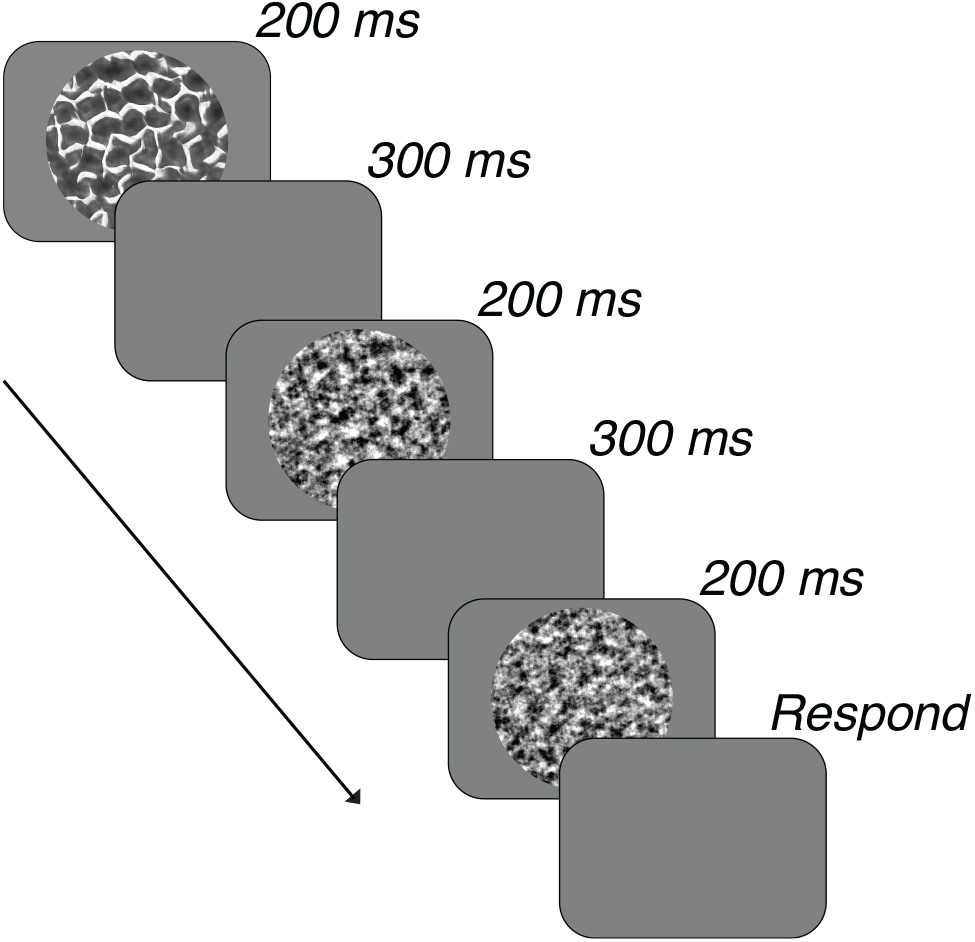
Task structure. Three images were flashed in quick succession. Observers were tasked with judging whether the first two images or the last two images came from the same category.

Trials were organized in blocks in which the naturalness level of the texture images could vary between 0.02 and 1. Each trial’s one or two naturalistic textures were presented at an equal level of naturalness. Trials were organized in two randomly interleaved staircases (1 one-up/two-down, 1 one-up/three-down) initialized to naturalness=0.23 (Equation 1) after 3 practice trials at full naturalness. Staircases ran for 160 trials (80/staircase) within each block. Blocks were run for a single texture family at a time, and for a single condition in each experiment (e.g., each blur level or each scaled size). Staircase methods were only used to distribute trials at naturalness levels close to threshold; the final convergence levels of the staircases were not used in further analyses.

For each block of trials, we maximized the likelihood of a logistic function fit to the psychometric data. The function was parameterized with a threshold, slope and lapse rate (Wichmann & Hill, 2001a). Naturalness sensitivity was defined as the reciprocal of the measured threshold. To compute error estimates of the sensitivity we used a parametric bootstrap method (Wichmann & Hill, 2001b) in which we repeatedly resampled and refit data from the best fit sigmoid function to get a bootstrap distribution of sensitivities. Error bars on sensitivities are reported as the 2.5% and 97.5% percentiles of the bootstrap distribution.

## Experimental results

Here, we report the results of a series of experiments to measure the ability of human observers to discriminate images of naturalistic texture from spectrally-matched noise. We used the Portilla & Simoncelli texture synthesis method to generate a continuum of images that smoothly vary in the strength of their naturalistic structure. Experiments using these images allowed us to quantify the “naturalness sensitivity” of observers in response to four different experimental manipulations. In experiment #1 we evaluated the role of high retinal spatial frequencies in naturalness sensitivity by blurring the texture images. In experiment #2 we tested the scale-invariance of naturalness sensitivity by rescaling images to different sizes, which approximates the effect of moving images towards or away from observers. In experiment #3 we used the results of the first two experiments to predict observer sensitivity to images that have been both rescaled and blurred. Finally, in experiment #4 we measured naturalness sensitivity for images presented in the peripheral visual field.

### Experiment #1: Image blur

We first measured how naturalness sensitivity changes with image blur. To blur images, we used a range of low-pass filters differing in corner frequency. Specifically, the original texture images were converted to their two-dimensional Fourier spectra, then multiplied by a low-pass filter. Each low-pass filter was defined by a raised-cosine edge that went from full to zero gain over the span of half an octave, with its corner frequency defined as the point of 50% gain. Applying the inverse Fourier transform created the final, filtered image. All filtering was done prior to image vignetting. Qualitatively, as images became more strongly blurred it became harder to distinguish texture images from noise images (Figure 4).

**Figure 4.**
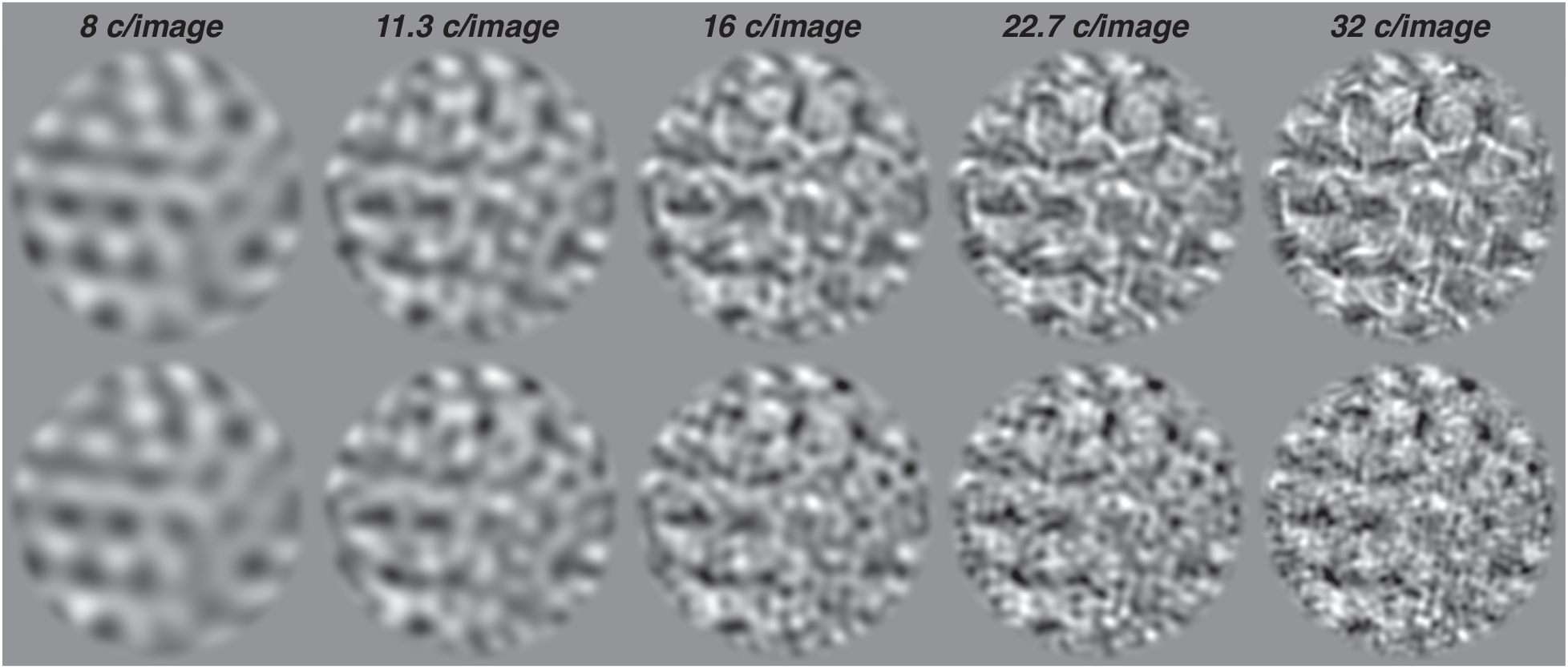
Example images for blur experiment. Examples of naturalistic (top, naturalness=0.376) and noise (bottom) images that have been low-pass filtered. From left to right, object spatial frequencies have been removed that are higher than 8, 11.3, 16, 22.7, and 32 c/image (images are 320 pixels wide, so this corresponds to spatial period cutoffs of 40, 28.3, 20, 14.1, and 10 pixels respectively). These images were presented at a distance of 1.42 m, such that their diameter measured 4 degrees of visual angle. At that size, these corner frequencies correspond to retinal spatial frequencies of 2, 2.8, 4, 5.7, and 8 c/deg.

For all 3 observers and 5 texture families, sensitivity rapidly improved with increasing corner spatial frequency up to approximately 11 c/deg, beyond which sensitivity saturated (Figure 5). There are two possible reasons for this saturation. One possibility is that the visual system only uses retinal spatial frequencies up to a corner frequency of 11 cycles per degree when performing this task, effectively representing a *texture acuity limit* (*f_acuity_*) for naturalness perception. The other possibility is that the texture images themselves contain no task-relevant information above this frequency. The texture synthesis algorithm generates images with no task-relevant information above object spatial frequencies of 113 cycles per image (see Methods, Image generation), a frequency that we refer to as the *synthesis limit* (*f_synth_*). Accordingly, when images are presented at a size of 4 degrees (as in this experiment) this synthesis limit corresponds to a retinal spatial frequency of 113/4 = ^~^28 c/deg. Our empirical measurement of the texture acuity limit, 11 c/deg, is well below this synthesis limit. This suggests that the measured texture acuity limit represents a feature of the visual system, rather than a feature of the texture images.

**Figure 5.**
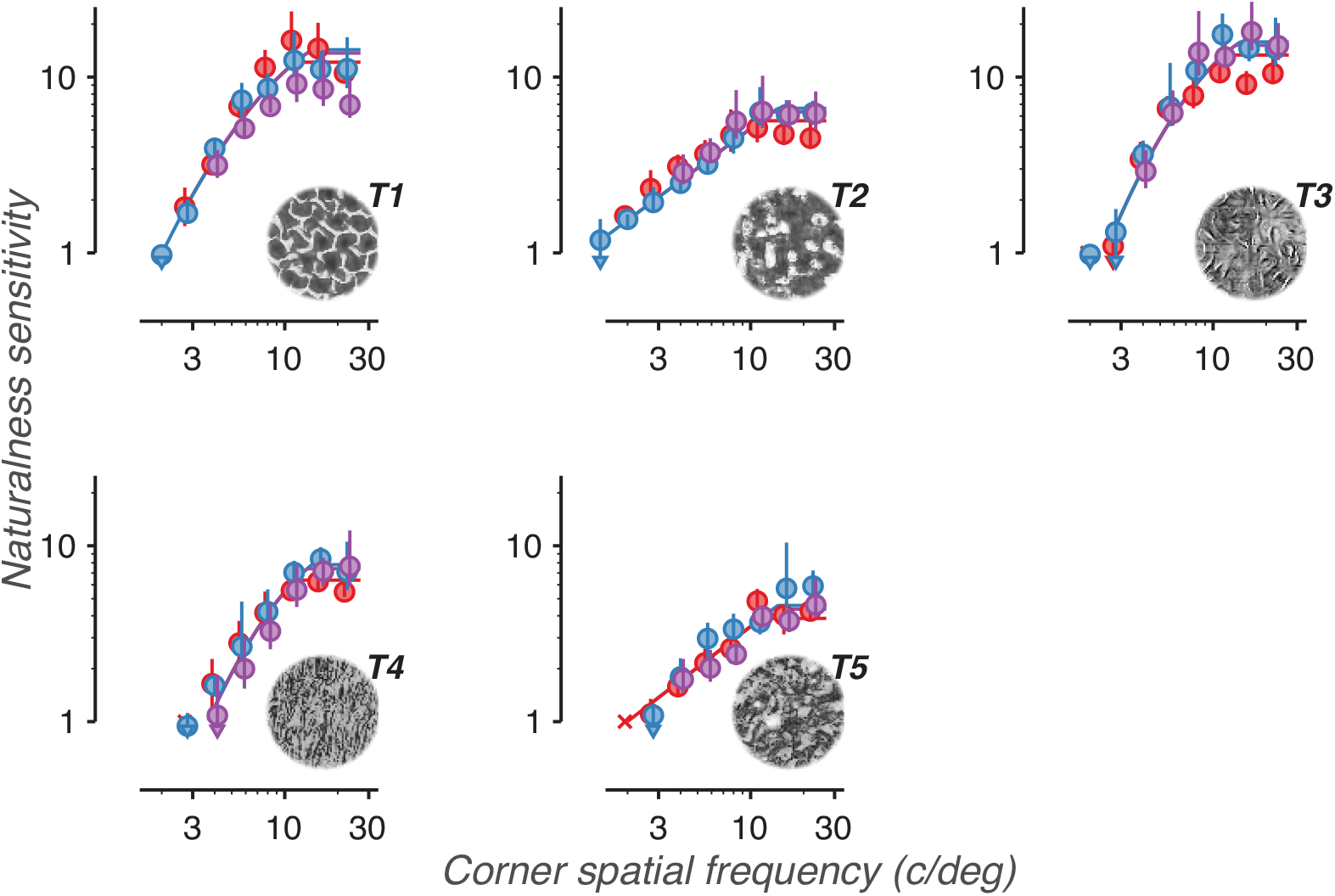
Naturalness sensitivity is reduced by image blur. Naturalness sensitivity vs. corner retinal spatial frequency for 5 different texture families. Data points labeled X denote conditions that were too difficult to produce a stable level of sensitivity. Colors distinguish the 3 observers. Error bars represent bootstrapped 95% confidence intervals. Arrows denote conditions in which the confidence intervals fell below the bottom edge of the plot. Solid lines denote descriptive fits based on data from all four experiments.

### Experiment #2: Image rescaling

By itself, the results of the blur experiment cannot disambiguate whether naturalness sensitivity is set by properties of the image set (the informativeness of different object spatial frequencies) or properties of the visual system (the filtering of different retinal spatial frequencies). An experimental manipulation that can decouple these two factors is changing viewing distance. As the distance between the image and the observer decreases, fine details in the image, which exist in high object spatial frequencies, shift into lower retinal spatial frequencies and become easier to see. If naturalness sensitivity is primarily set by a texture acuity limit then decreasing viewing distance should lead to higher sensitivity.

For this experiment we rescaled texture and noise images to larger and smaller sizes, a process that approximates physically placing it closer or further away from the observer. To achieve this effect, we resized images (*imresize(*) in Matlab, bicubic interpolation), to produce a range of image diameters from 40 pixels (0.5 degrees) to 640 pixels (8 degrees). To achieve rescaled sizes of 16 and 32 degrees, we presented the 640 pixel images at viewing distances of 71 and 35.5 cm, respectively. Qualitatively, naturalistic structure was easier to perceive at larger sizes (Figure 6).

**Figure 6.**
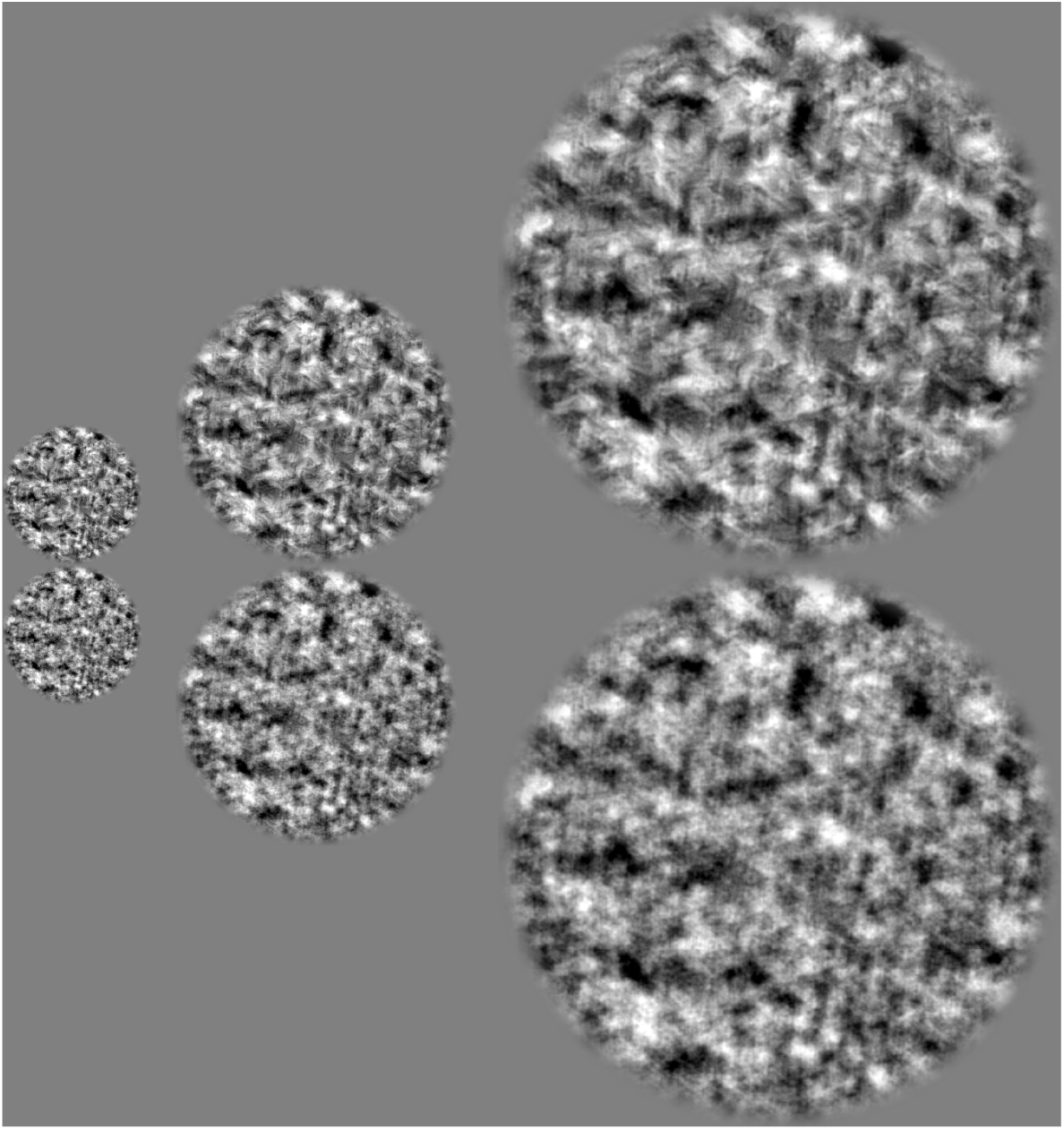
Example images for scaled size experiment. Naturalistic (top, naturalness=0.087) and noise (bottom) images are rescaled to diameters of 160, 320, and 640 pixels. When presented in the experiment, these images took up 2, 4, and 8 degrees of visual space, respectively.

The texture images contain no task-relevant information above the synthesis limit *f_synth_*=113 c/image. When images are presented at larger scaled sizes, the corresponding retinal spatial frequency of the synthesis limit decreases. For image sizes of 4, 8, and 16 degrees, the synthesis retinal frequency falls from 28 c/deg, to 14 c/deg, to 7 c/deg. Accordingly, there exists a scaled image size sufficiently large that the synthesis limit (a property of the image) will fall below the texture acuity limit (a property of the visual system). At this size, the only image features that would still lie above the texture acuity limit are those that contain no task-relevant information and increasing the size of the image would no longer improve naturalness sensitivity. The texture acuity limit thus predicts that saturation should begin at a size equal to the ratio of the synthesis limit to the texture acuity limit:

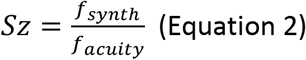

The texture acuity limit of 11 c/deg measured in Experiment #1 predicts a saturation size of about 10 degrees.

For all 3 observers and 5 textures, we found that sensitivity increased with scaled size up to approximately 8 degrees, after which sensitivity saturated (Figure 7). The observed point of saturation corresponds closely to the texture acuity limit prediction of 10 degrees. These results are consistent with naturalness sensitivity being limited by a single texture acuity limit, a property of the visual system, that applies to both experiments #1 and #2. Furthermore, note that in both experiments textures 2 and 4 qualitatively show a noticeably shallower slope than textures 1, 3, and 5. These two observations suggest that texture acuity limits (features of individual observers) and sensitivity curve shapes (features of individual texture families) may be applied across distinct experimental conditions.

**Figure 7.**
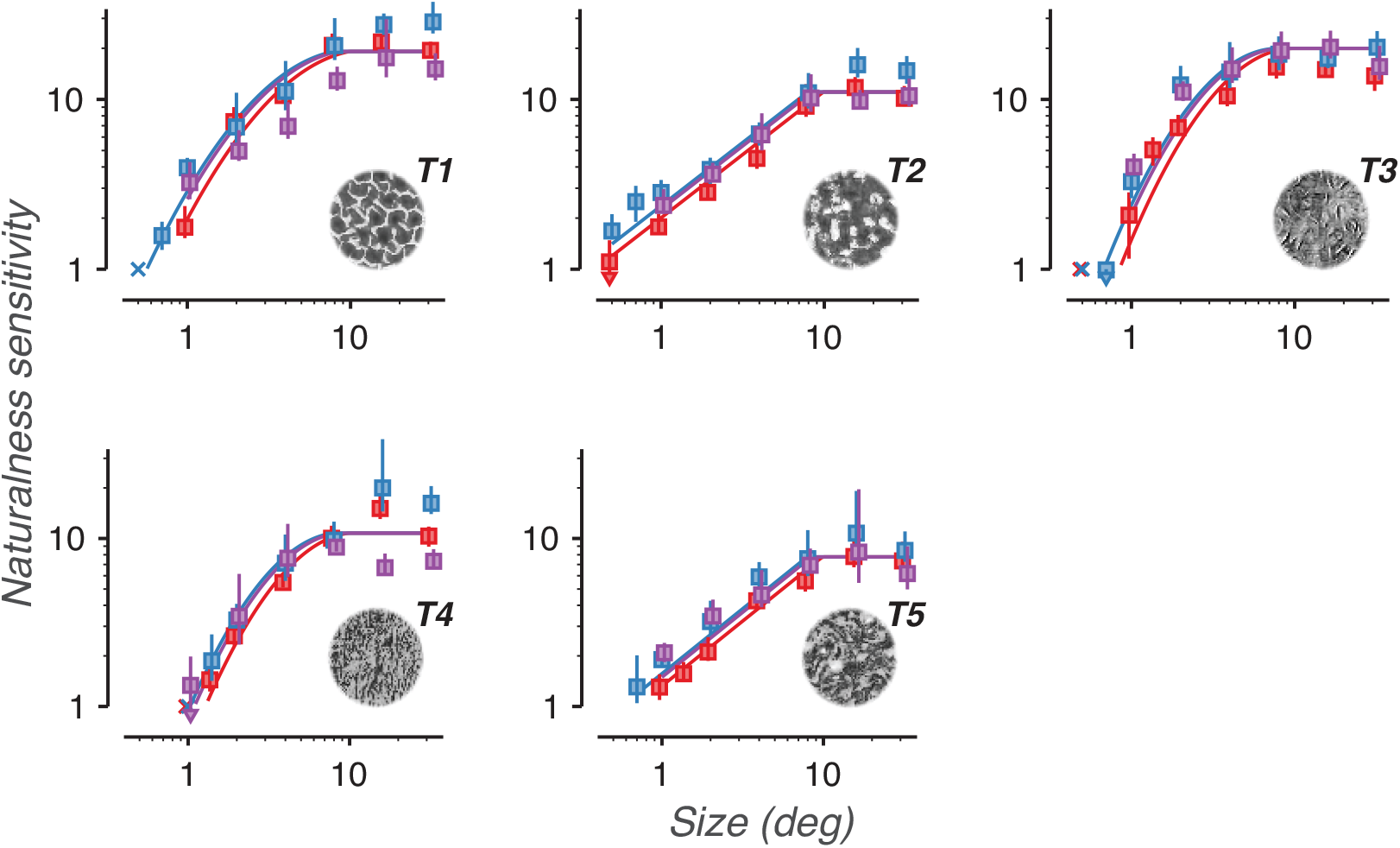
Naturalness sensitivity increases as textures are scaled to larger sizes. Naturalness sensitivity vs. scaled image size for 5 different texture families. Data points labeled X denote conditions that were too difficult to produce a stable level of sensitivity. Colors distinguish the 3 observers. Error bars represent bootstrapped 95% confidence intervals. Arrows denote conditions in which the confidence intervals fell below the bottom edge of the plot. Solid lines denote descriptive fits based on data from all four experiments.

### Experiment #3: Resized, blurred textures

The observed relationship between scaled size and sensitivity relies on the distribution of task-relevant information in different object spatial frequencies. Blurring the images has the effect of removing information in high object spatial frequencies. That is, blurring effectively lowers the synthesis limit of the image set. The results of the blurring and scaling experiments predict that lowering the synthesis limit should lead to saturation at a smaller scaled size. To test this interaction, we generated blurred, rescaled images by first low-pass filtering the images at a corner frequency of 22.6 cycles/image, then rescaling them as in Experiment #1. This amount of blur corresponds to the 4^th^ pair of images from the left in Figure 4. Substituting 22.6 c/image into Equation 2 predicts that sensitivity should saturate for sizes greater than 2 degrees. For 2 observers and all 5 textures, we found that the measured saturation size closely matched this prediction (Figure 8).

**Figure 8.**
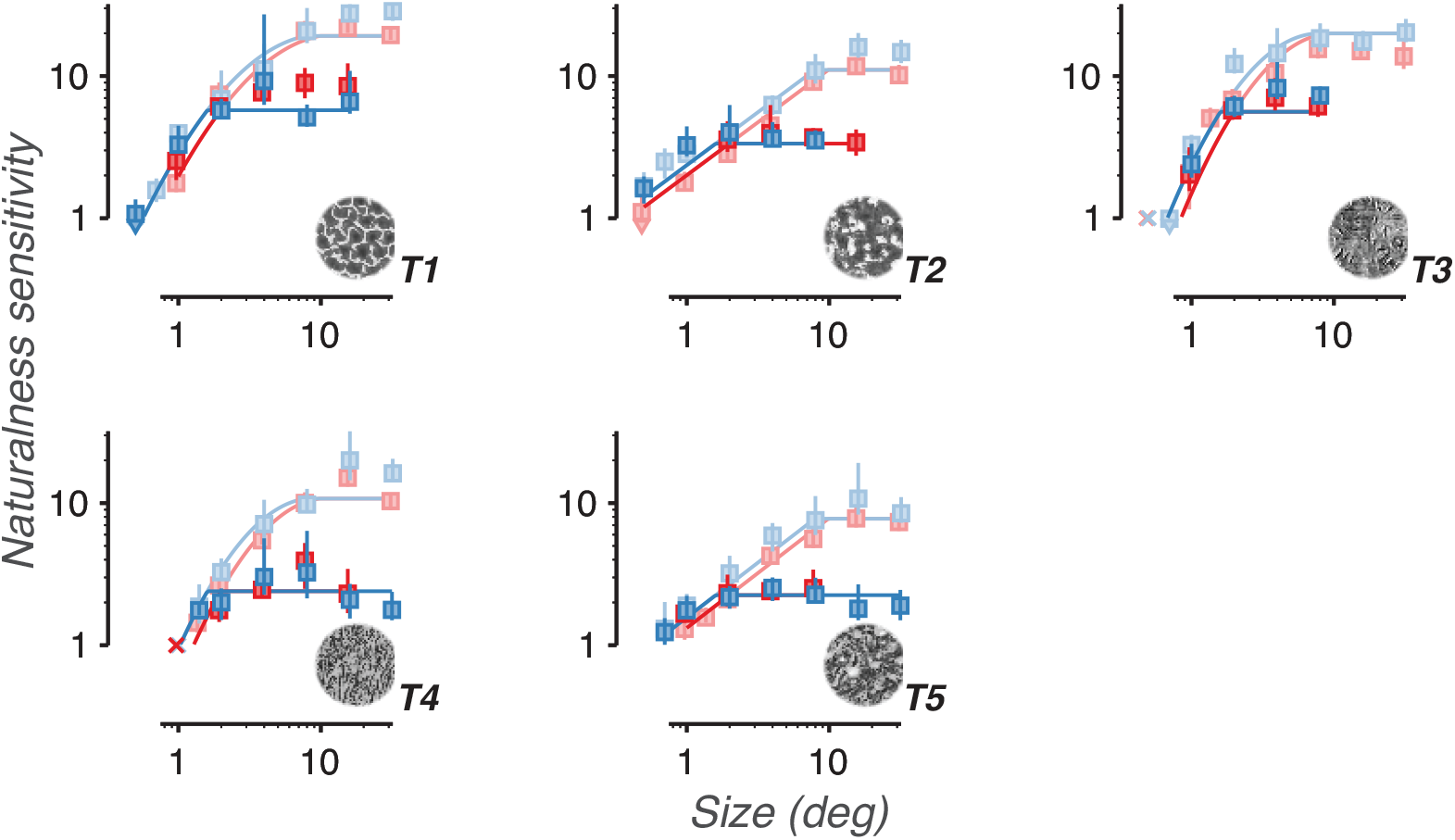
Naturalness sensitivity saturates at smaller scaled sizes for blurred images than for unfiltered images. Naturalness sensitivity vs. scaled image size for 5 different texture families. Light points and lines represent unfiltered images (data replotted from Figure 7). Dark points and lines represent images low-pass filtered at a corner frequency of 22.6 c/image. Data points labeled X denote conditions that were too difficult to produce a stable level of sensitivity. Colors distinguish two observers. Error bars represent bootstrapped 95% confidence intervals. Arrows denote conditions in which the confidence intervals fell below the bottom edge of the plot. Solid lines denote descriptive fits based on data from across all four experiments.

This experiment demonstrates that, for low-pass filtered images, naturalness sensitivity is invariant to changes in scale over an 8:1 increase in image size (2 degrees to 32 degrees). This scaling of image size corresponds to a shift in the retinal spatial frequencies of the image over three octaves (e.g., from 11.3 to 1.4 c/deg). This scale-invariance demonstrates that observers appear to be equally effective at extracting task-relevant information located in either low or high retinal spatial frequencies. We conclude that the reliance of naturalness sensitivity on high-frequencies is not due to superior visual processing of image features in high *retinal* spatial frequencies. Rather, these measurements suggest that the texture images may be intrinsically more informative in high *object* spatial frequencies. We will address this question more quantitatively in the next section (“Descriptive fit”) where we describe a model observer analysis of the images.

### Experiment #4: Images in the visual periphery

Visual acuity peaks in the center of gaze and steadily drops at locations further eccentric. To characterize if and how quickly texture acuity declines with visual eccentricity, we presented texture images in the peripheral visual field.

Images were presented at rightwards horizontal displacements from the point of fixation, measured as the distance from the center of the fixation point to the center of the image. Thus, when the 4 degree diameter image was presented at an eccentricity of 8 degrees, it spanned eccentric locations between 6 and 10 degrees. To achieve eccentric locations larger than 12 degrees, the monitor was moved to a distance of 71 cm and images were rescaled to a diameter of 160 pixels. For this experiment, we recorded eye movements using an EyeLink eye tracker and verified eye position stability. Subjects received auditory feedback on whether they maintained stable fixation throughout the trial. Qualitatively, as images are presented further peripherally, texture becomes more difficult to distinguish from noise (Figure 9).

**Figure 9.**
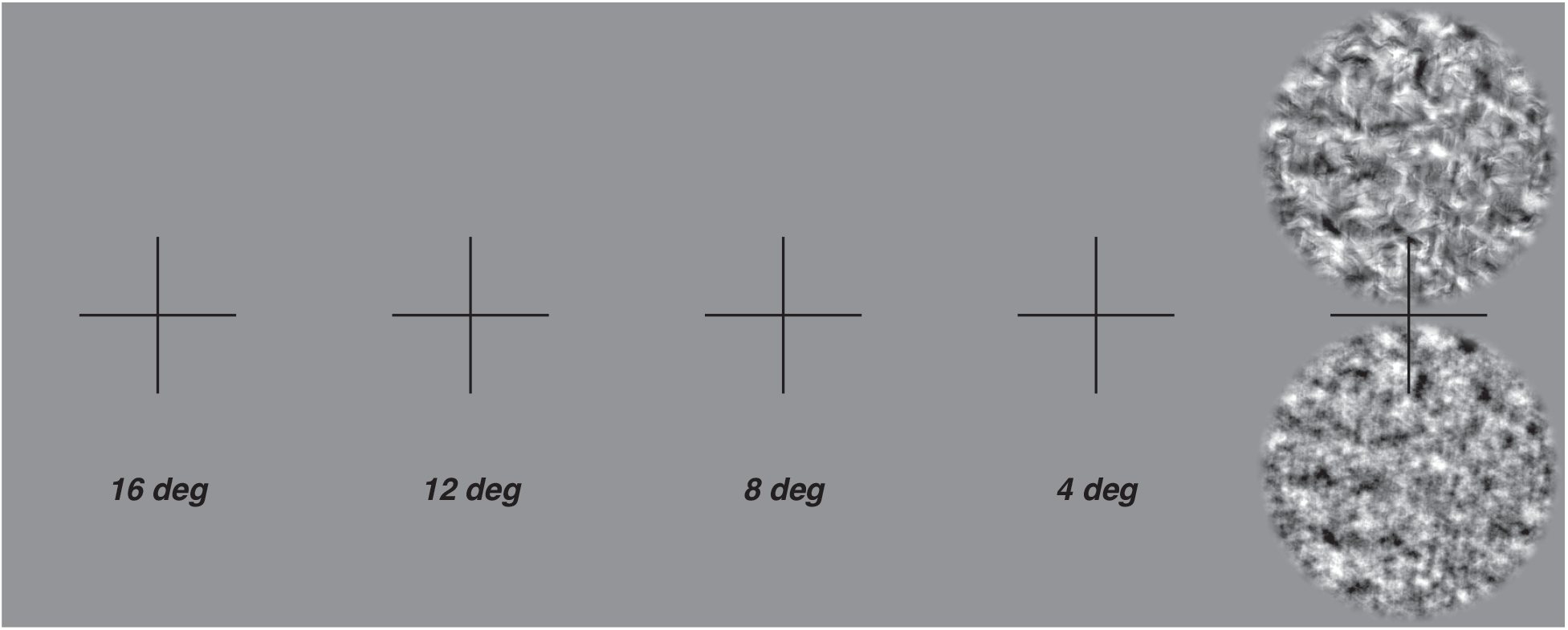
Demonstration of visual periphery experiment. Hold the above figure at about arm’s length, such that the black cross takes up ^~^2 degrees of visual angle (approximately the width of your thumb at arm’s length). Fixating at the center of each cross will present the naturalistic (naturalness = 0.231) and noise images at a diameter of 4 degrees, at an approximate distance from fixation of 16, 12, 8, 4, and 0 degrees, from left to right. Note that this demonstration (in which two images are presented at the same time) is not a reproduction of the experimental setup (in which only one image was flashed at a time). As the images come closer to the center of gaze, it becomes easier to discriminate texture from noise.

For all 3 observers and 5 texture families, sensitivity smoothly declined with increasing eccentricity (Figure 10). Similar to experiments 1 & 2, texture families 2 and 5 qualitatively demonstrate a shallower decline than texture families 1, 3, and 4. This suggests that these differences in slope may be related to texture family-specific distributions of information over different object spatial frequencies. Accordingly, texture acuity may be an observer-specific property of the visual system that is applied equally to all texture families. The slopes in Figure 10 rely on both of these factors: the distribution of information in each texture set and how texture acuity falls with eccentricity. Extracting how texture acuity falls with eccentricity requires a model that can disentangle these two factors.

**Figure 10.**
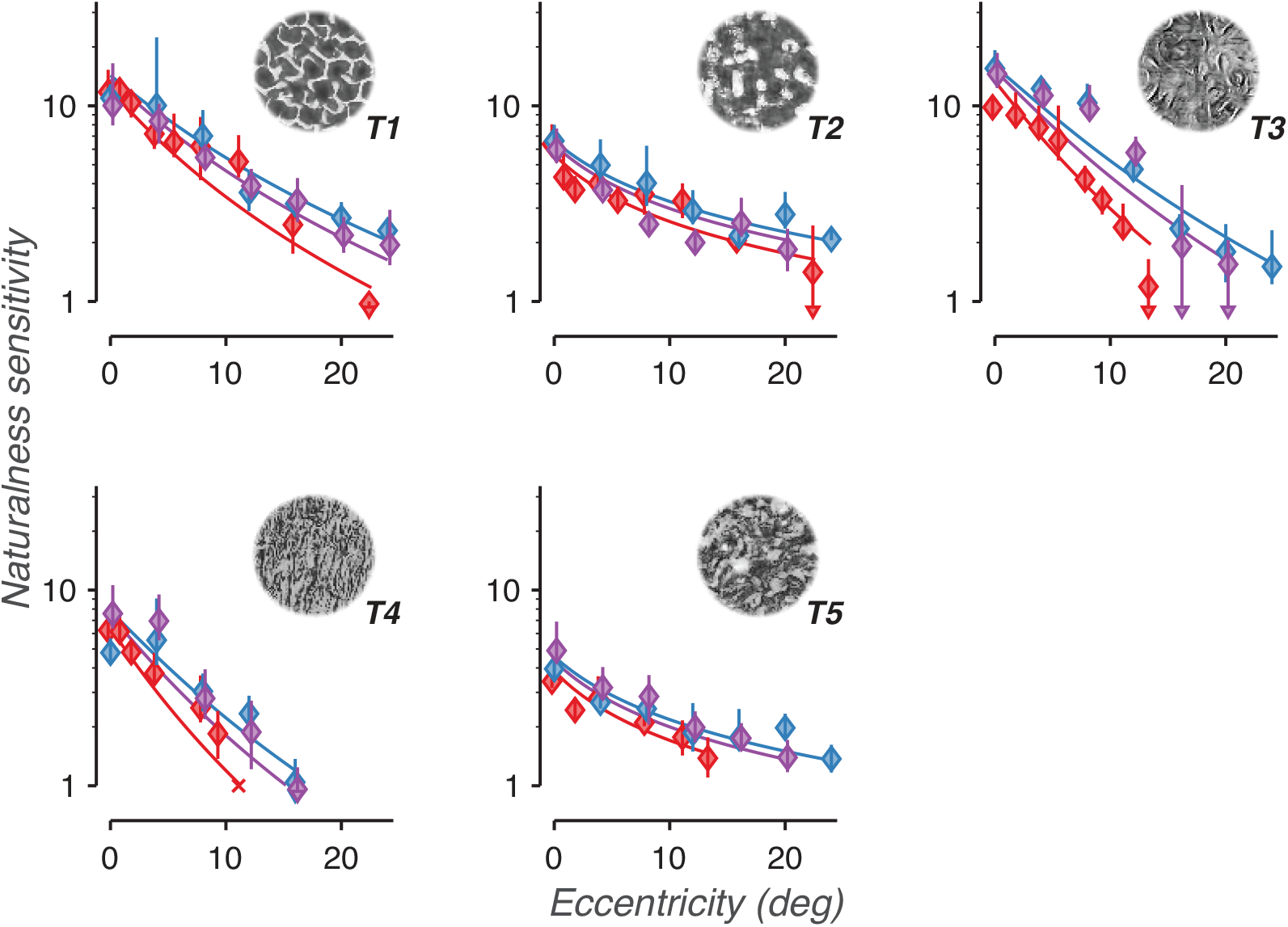
Naturalness sensitivity is lower in the peripheral visual field. Naturalness sensitivity vs. presentation location in the visual field for 5 different texture families. Data points labeled X denote conditions that were too difficult to produce a stable level of sensitivity. Colors distinguish the 3 observers. Error bars represent bootstrapped 95% confidence intervals. Arrows denote conditions in which the confidence intervals fell below the bottom edge of the plot. Solid lines denote descriptive fits based on data from all four experiments.

### Experiment #5: Grating acuity

In our last experiment, we sought to compare texture acuity in the visual periphery to measurements from a more established task that is known to be limited by contrast (and thus depend on early visual mechanisms). Observers performed a coarse orientation discrimination task on images of sinusoidal gratings presented at different locations in the visual periphery. We used staircase procedures to change the retinal spatial frequency of the grating in order to find their *grating acuity*: the highest retinal spatial frequency at which observers could still consistently complete the task.

Observers used the same match-to-sample task structure as the naturalness task. Three apertured sinusoidal gratings (4 degree diameter circle) were flashed, and the first and last grating differed in orientation by 90 degrees. The middle grating could match either the first or the last grating in orientation. Observers reported whether the first two or second two gratings matched in orientation. The retinal spatial frequency of all three gratings was the same, and changed trial-to-trial using the same staircase procedure as in the naturalness experiments (1 to 38.1 c/deg, 0.25 octave spacing). In different blocks of the experiment, gratings were presented at different locations in the peripheral visual field, using the same methods as experiment #4.

The results of Experiment #5 are plotted in Figure 12. Qualitatively, observers reported that threshold spatial frequency corresponded to the point at which gratings were barely visible (Watson & Robson, 1981). We will postpone further discussion of these results until we have extracted comparable measurements of texture acuity from the descriptive model detailed in the next section.

## Analysis

We have described naturalness sensitivity measurements from four distinct experimental conditions. Four aspects of our psychophysical results suggest that a common mechanism underlies these results. First, in all cases, sensitivity declines as high spatial frequency image features are attenuated or made more difficult to see. Second, the shapes of these declines are idiosyncratic to each texture family. Third, when a texture family exhibited a particular shape in one experiment, it tended to exhibit that same shape in all other experiments. Finally, the limits of naturalness perception seem to be set by two distinct spatial frequencies. The first is a *synthesis limit*: an object frequency (measured in cycles per image) set by the texture synthesis procedure. The second is a *texture acuity limit*: a retinal frequency (measured in cycles per degree) set by the limits of the visual system. These observations set the form of a simple, descriptive model of naturalness sensitivity that we use to fit all experimental data simultaneously.

The model contains two cascaded stages. The first stage fits, for each texture family, a *cumulative sensitivity function*: a simple relationship between naturalness sensitivity and the corner object spatial frequency of a low-pass filter (Figure 11A / Figure 13). This stage estimates how well human observers extract information from the accessible object spatial frequencies of an image set, and its fit can be directly compared to the naturalness sensitivity of a model observer (see Appendix). This stage uses a single exponentiated quadratic function per texture family (Equation 6, three parameters/family, 15 total) that is shared across all experimental conditions and observers.

**Figure 11.**
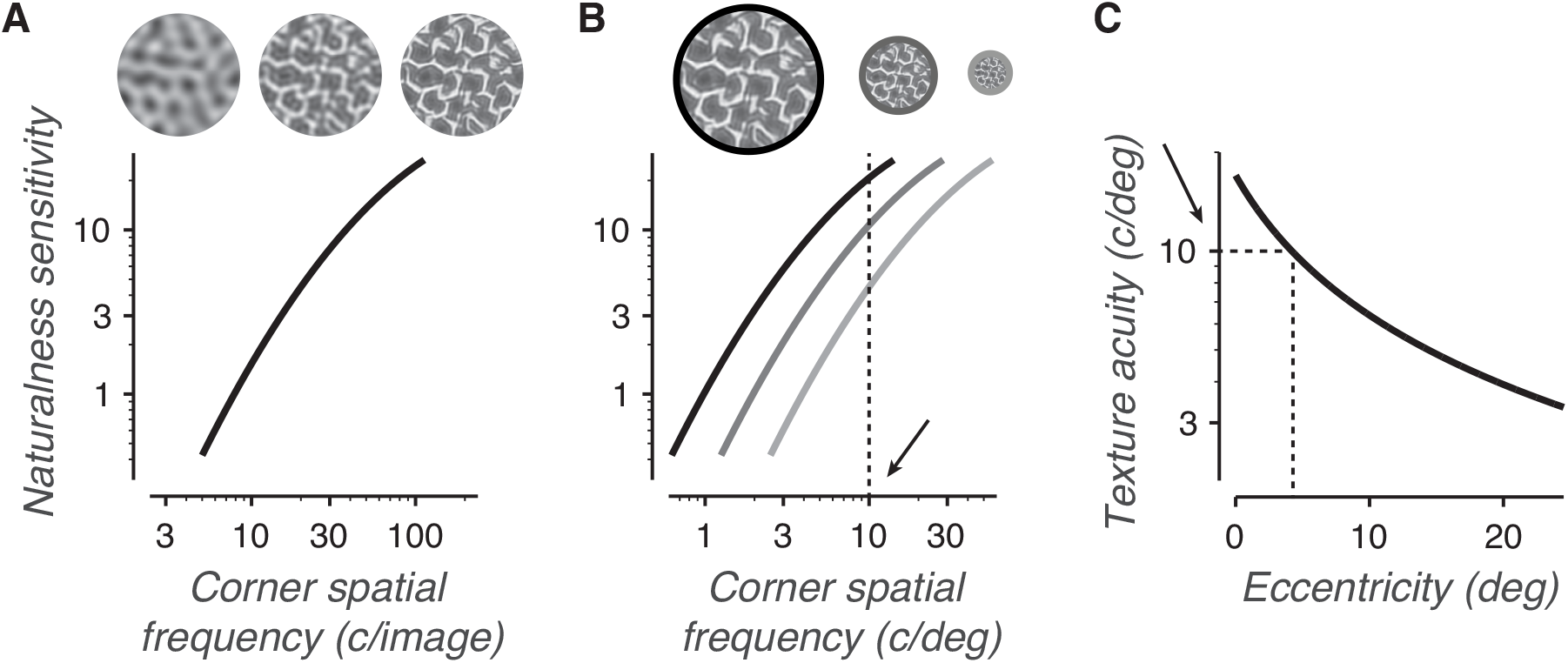
A descriptive model for naturalness sensitivity measurements. A) For each texture family, we fit a cumulative sensitivity function that maps how naturalness sensitivity improves as observers gain access to features in increasingly high object spatial frequencies. The function plots sensitivity vs. the corner frequency of a low-pass filter. As more high frequencies are filtered out of the image (moving right to left on the plot), sensitivity declines. B) In the experiment, texture images are presented at a scaled size measured in degrees of visual angle. This size determines the mapping of the cumulative sensitivity function from units of object spatial frequency (cycles/image) to units of retinal spatial frequency (cycles/degree). The black, dark gray, and light gray curves correspond to the sensitivity curve in A) shifted for presentation sizes of 8, 4, and 2 degrees, respectively. The intersections of these curves with a vertical line denotes how sensitivity changes with size for a specific value of the texture acuity limit. Moving further into the visual periphery moves the vertical line left. The dotted line corresponds to a texture acuity limit of 10 c/deg. C) Texture acuity is highest at the center of gaze and lower in the peripheral visual field. The texture acuity function describes, for each observer, how texture acuity falls with eccentricity. The dotted lines illustrate that when images are presented at 4 degrees eccentricity, texture acuity is 10 c/deg.

**Figure 12.**
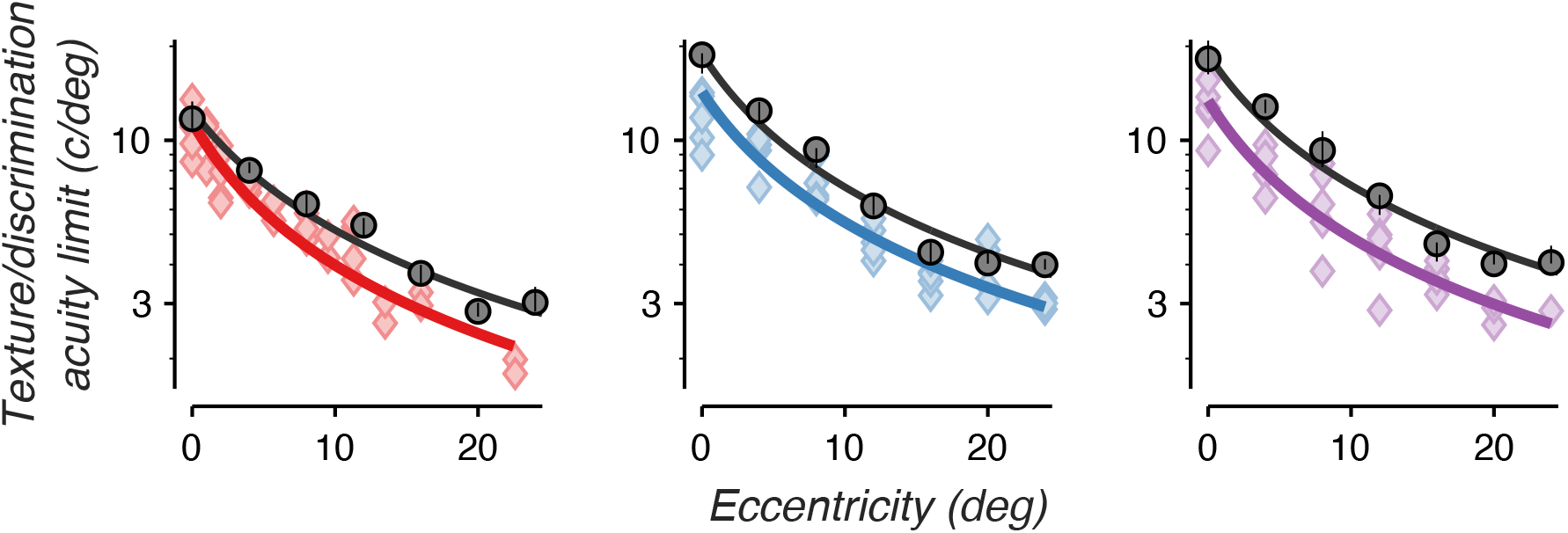
Texture acuity and orientation discrimination acuity show similar rates of decline with eccentricity. The colored lines plot descriptive fits of texture acuity vs. eccentricity in the visual field. Individual scatter points denote texture acuity limits predicted from individual conditions in Experiment 4, mapped using each texture family’s cumulative sensitivity function (Figure 13). Each plot denotes a separate observer, with colors denoting specific observers as in previous figures. The black points denote grating acuity measurements from an orientation discrimination task (90 degree orientation difference, grating RMS contrast = 10%). The black curves are maximum likelihood fits of the grating acuity function.

The second stage of the model then “presents” the texture family at a given scaled size and location in the visual field (Figure 11B-C), determines the relevant corner frequency (synthesis limit or texture acuity limit), and computes the resulting naturalness sensitivity of the observer. This stage uses a single function per observer (Equation 4, two parameters/observer, 6 total) that describes how texture acuity falls from the center of gaze to the peripheral visual field (Figure 11C / Figure 12) and is shared across all experimental conditions and texture families.

### Analysis methods

We assume that, in all four experiments, naturalness sensitivity depends solely on the highest object spatial frequency available to the observer. This controlling corner spatial frequency in each experimental condition can be set by one of two factors. The first is an intrinsic object spatial frequency, the image limit:

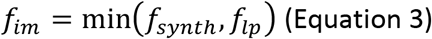

that is either a property of the texture synthesis process (*f_synth_* = 113 cycles/image, see Experimental methods: Image generation) for unfiltered conditions, or a result of low-pass filtering for blur conditions (*f_lp_*). This parameter represents the spatial frequency above which the image no longer contains task-relevant information. The units of this limit are in cycles/image.

The second factor is a property of the visual system, the texture acuity limit, which describes the highest retinal spatial frequency that the observer can use to solve the task. We assume that, like contrast detection acuity (Anderson et al., 1991; Robson & Graham, 1981), texture acuity has a finite value at the center of gaze and falls inversely with eccentricity:

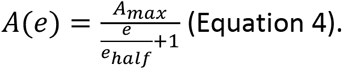

*A_max_* is a fit parameter for the foveal texture acuity limit, and should correspond to the measured point of saturation in Experiment #1. *e* is the visual field eccentricity at which the image was presented. *e_half_* is a fit parameter that corresponds to the visual field eccentricity at which texture acuity falls to half of the foveal limit, and is equivalent to the x-intercept measurement discussed in previous measurements of peripheral acuities (Levi & Klein, 1990; Virsu & Rovamo, 1979). This texture acuity function has units of cycles/degree.

Experimental conditions live in one of two regimes. In the first, observers can see all task-relevant information present in the image and the controlling spatial frequency is equivalent to the image limit. In the second, observers cannot use all task-relevant information present in the image because some of it lies beyond the texture acuity limit. In this second regime, the controlling spatial frequency is set by the texture acuity limit. We define a normalized controlling frequency *f_norm_* as:

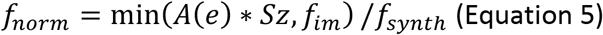

Where *Sz* is the scaled size of the image in degrees. Because acuity is always positive and *f_im_* is always less than or equal to *f_synth_*, *f_norm_* is bounded as 0 < *f_norm_* ≤ 1.

We observed that for some texture families, sensitivity began to saturate when using object spatial frequencies near the synthesis limit. To account for this saturation, we fit the cumulative sensitivity function as an exponentiated quadratic function:

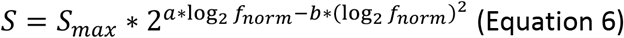

where *S_max_*, *a*, and *b* are fit parameters constrained to be positive. On a plot of log sensitivity versus log frequency (as in Figure 13) this function takes the shape of a quadratic curve. Because *f_norm_* ≤ 1, *S_max_* is the maximum possible naturalness sensitivity for each texture family. The cumulative sensitivity function is used to set the threshold parameter (*T* = 1/*S*) of a logistic function that describes how performance varies with naturalness.

**Figure 13.**
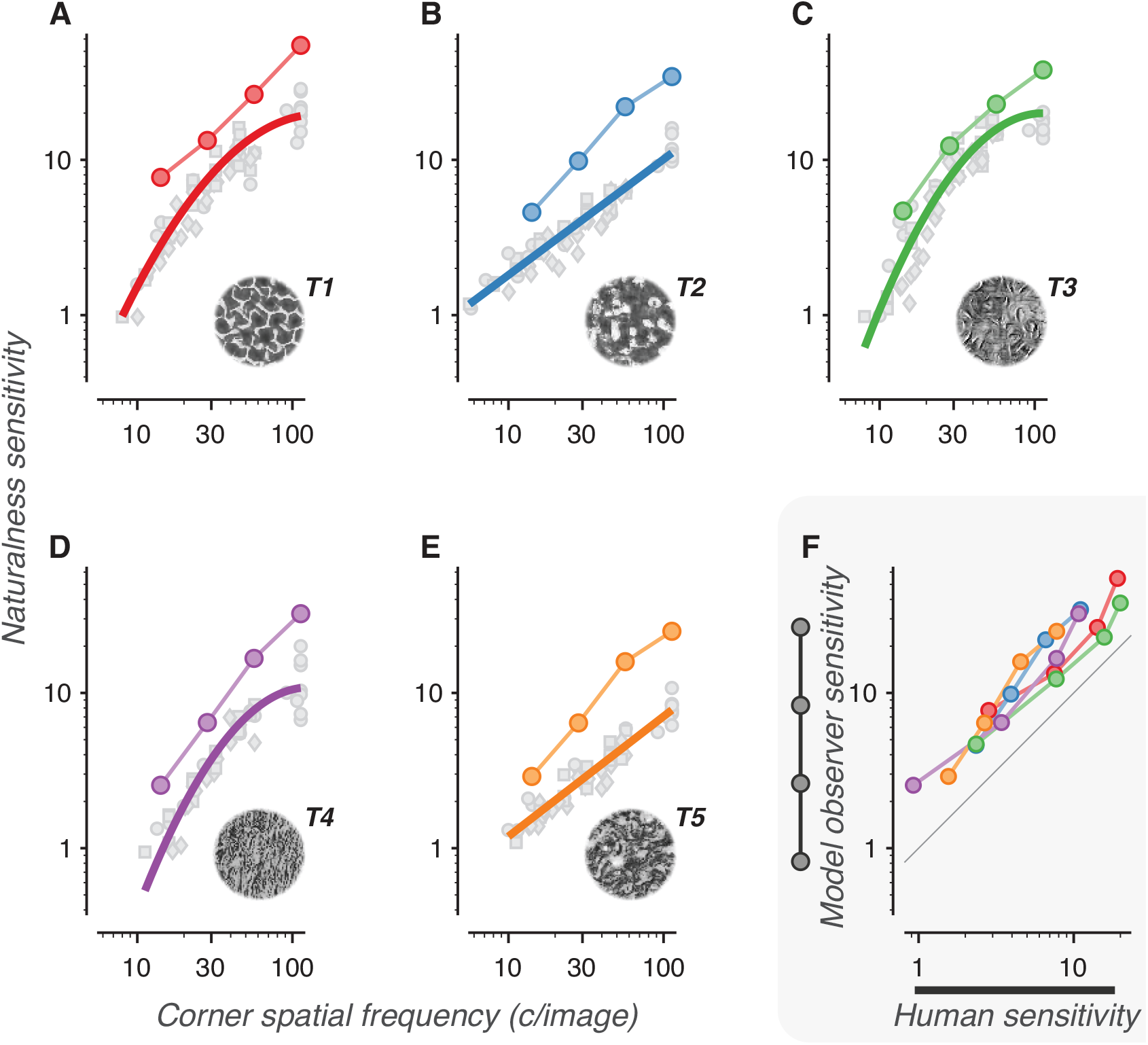
Cumulative sensitivity function fits and model observer sensitivities for each texture family. A-E) Cumulative sensitivity functions (best fit curves of naturalness sensitivity vs. low-pass corner frequency) are plotted as thick colored lines for each of the 5 texture families. Scatter points denote individual conditions from all 4 experiments, where each condition’s corner spatial frequency has been calculated based on its low-pass filtering, size, and presentation eccentricity. Different symbols represent data points from different experiments (squares - rescaling, circles – blur, diamonds – visual periphery). The naturalness sensitivity of the model observer is plotted against low-pass corner frequency as colored circles connected by a thin line. F) Model observer sensitivities (data points from the colored circles in A-E, thin lines) are plotted against descriptive fit sensitivity measurements (measured at 4 points along the colored thick lines in A-E). Colors denote individual texture families, as in A-E. The black line denotes unity.

Using these functions, we found the parameter set that maximized the likelihood of the fit for all individual trials. In its final form, the model fits 23 parameters to more than 71,000 trials distributed over 20 distinct experimental conditions for the 3 observers (55 curves total). The 3 sensitivity curve parameters from Equation 6 are fit individually for each of the 5 texture families (15 parameters total). The 2 acuity parameters from Equation 4 are fit for each observer (6 parameters total). The logistic function slope and lapse rate parameters are shared among all observers (2 parameters).

Grating discrimination acuity curves (Figure 12, black curves) were separately fit to the data from Experiment #5 in a manner similar to that used for texture acuity. For each of the three observers, we used the acuity formula (Equation 4) to determine thresholds of the sigmoid function at each eccentricity, then maximized the likelihood of the fit.

### Analysis results

The results of the descriptive fit are summarized in Figures 12 and 13. Figure 12 plots fits of each observer’s texture acuity against visual eccentricity. Texture acuity is highest at the fovea (13.04 c/deg average, ± 1.43 std. dev.) and declines inversely proportional to eccentricity. Texture acuity fell to half of its maximum value at an eccentricity of 5.75 degrees (average ± 0.44 std. dev.). We compared this rate of decline to that of grating acuity, plotted as the set of black curves in Figure 12. Note that for this task, the absolute acuity values depend on grating contrast; grating acuity shifts up or down with increased or decreased image contrast. Nonetheless, we expect that the relative rate of decline in grating acuity with eccentricity should be the same when using different contrasts (Anderson et al., 1991; Robson & Graham, 1981). Grating acuity fell to half of its maximum at an eccentricity of 6.40 degrees (average ± 0.57 std. dev.), a rate of decline very similar to that of texture acuity.

Figure 13A-E plots cumulative sensitivity functions (fits of naturalness sensitivity vs corner frequency) for each texture family (thick lines). Plotted on the same graphs are the sensitivities and estimated corner spatial frequencies for all individual experimental conditions that used that texture family (gray points). Their close correspondence suggests a parsimonious account of the data.

Though there is some heterogeneity among different texture families, all five plots show a rapid rise in sensitivity with corner frequency. The slope on these log-log plots corresponds to the exponent of a power function fit of sensitivity versus frequency. The average slope across the texture families is 1.06 (+- 0.297 std. dev., measured between 10 and 100 c/image), which corresponds to a linear relationship between sensitivity and frequency. In other words, each addition of an octave-wide frequency band to the texture images leads, on average, to a doubling in naturalness sensitivity.

### Efficiency

The cumulative sensitivity functions do not disambiguate whether 1) image features in high object spatial frequencies are intrinsically more informative than those in low frequencies or 2) observer processing is more efficient in high object spatial frequencies. To quantify the extent to which image features set the limits of naturalness discrimination, we built a model observer that measures the task-relevance of the texture statistics within each set of apertured images. Image aperturing led to a small amount of drift in each image’s texture statistics, such that different images synthesized to have the same naturalness could differ slightly in their statistics (Ziemba et al., 2018; Ziemba & Simoncelli, 2021). The model observer’s sensitivity corresponds to the naturalness level at which texture statistics are no longer a reliable cue for discrimination. The full details of the model observer’s implementation can be found in the Appendix.

We measured how the model observer’s naturalness sensitivity changed as we removed high object spatial frequency bands from the model (a process comparable to low-pass filtering the images). The end result is, for each texture family, a plot of the model’s naturalness sensitivity versus low-pass filter corner frequency. This is represented as the colored scatter points connected by a thin line in Figure 13A-E. For all texture families, the model observer’s sensitivity rapidly increases with corner frequency, approximately doubling with each addition of an octave-wide frequency band. This is the same average slope observed in the cumulative sensitivity functions.

We also measured the model observer’s sensitivity when it was limited to single frequency bands (a process comparable to bandpass filtering). Sensitivity was identical for bandpass and low-pass model observers when the single band of the bandpass model matched the highest band of the low-pass model. That is, the model observer’s sensitivity was fully set by the highest available object spatial frequency band, and the contribution of lower frequency bands was negligible.

We found that the sensitivity of the model observer covaries strongly with the sensitivity of human observers (Figure 13F, R^2^=0.91) and exhibits only slightly higher average sensitivity. The model observer partially captures the rank order of texture sensitivities, as well as the average slope of the sensitivity curve. The correspondence is not perfect, though. The model observer fails to capture particular idiosyncrasies of individual textures: the saturation at high object frequencies of textures 1, 3, and 4, and the shallower slope of textures 2 and 5. Nonetheless, we find that a model based purely on the image ensemble’s intrinsic uncertainty, with no added “neural” noise, closely tracks human performance on the texture task.

Task *efficiency* can be defined as the squared ratio of human sensitivity to model observer sensitivity (Barlow, 1978; Geisler, 2003). Observers have an average efficiency of 21.3% compared to the model observer (Figure 14). Efficiency is weakly bandpass, peaking at 26.5% at a spatial frequency of 28.3 c/image, and falling to 14.2% at the highest spatial frequency, 113.1 c/image. Note that, unlike other models, our model does not seek to be “ideal” in any sense. Our measurements of task efficiency should therefore be thought of as a comparison to an objective benchmark, rather than an absolute statement about a theoretical maximum.

**Figure 14.**
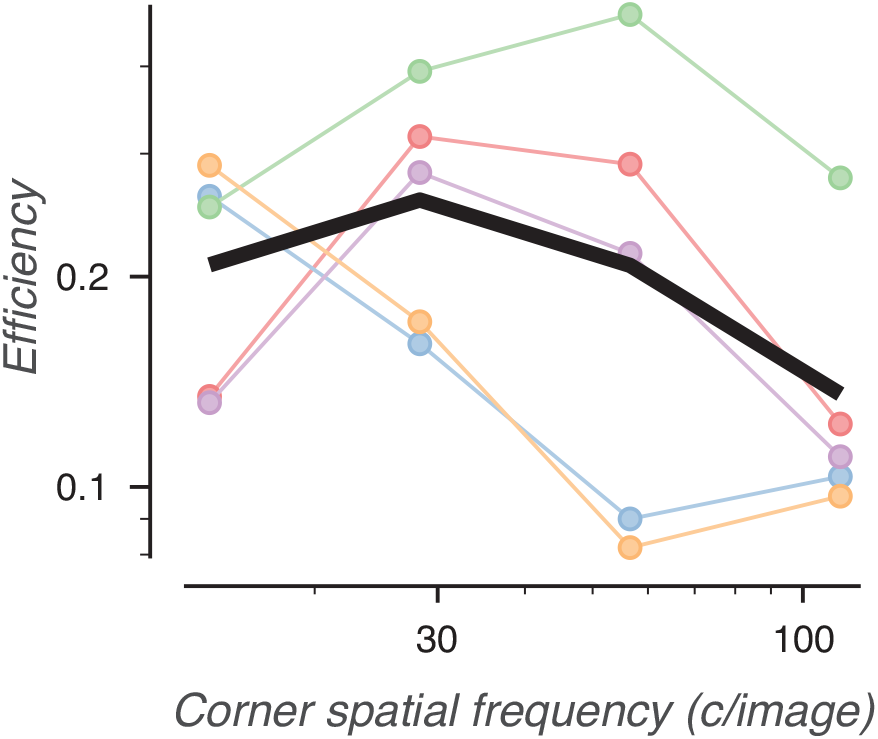
Efficiency of human naturalness sensitivity compared to the model observer. Efficiency is plotted against low-pass corner frequency. The black line is the average efficiency across textures. Colored points and lines represent the efficiencies of individual texture families, as in Figure 13.

#### To summarize

The texture image sets contain more task-relevant information in high object spatial frequencies than low frequencies. This rapid rise in information drives much higher human sensitivity when high frequencies are available, even after taking into account that observers are slightly less efficient at extracting information from high frequencies compared to low frequencies.

## Discussion

We have found that perceptual sensitivity to naturalistic texture stimuli relies on image features in high object spatial frequencies. As a result, sensitivity is primarily limited by a retinal spatial frequency, texture acuity, that varies across the visual field. A model observer analysis illustrates that high object spatial frequency image features carry more task-relevant information than low frequency features, and that human observers efficiently extract task-relevant information from all object spatial frequencies.

### Texture acuity limit

Observers performing the naturalness discrimination task cannot use image features that lie above the texture acuity limit. Similarly, observers cannot discriminate the orientation of two periodic gratings whose spatial frequencies lie above a grating acuity limit. Both texture acuity and grating acuity peak at the center of gaze and fall off with increasing eccentric distance. We find that both acuities decay at similar rates, falling to half of their foveal values at an eccentricity of ^~^6 degrees. It is parsimonious to assume that, due to the similarity in slope, both of these tasks are limited by contrast detection acuity. This decay is more gradual than some previously reported values for contrast acuity, which may be due to our relatively brief presentation time (Levi & Klein, 1990). Shorter presentation times and faster onsets degrade foveal acuity more strongly than peripheral acuity, and can accordingly lead to a shallower slope of acuity with eccentricity. Thus, we find that naturalness sensitivity, a task designed to isolate mid-level visual mechanisms, is nonetheless strongly limited by contrast acuity, an early visual mechanism.

### Scale-invariance

A visual task is considered *scale-invariant* if task performance stays consistent across changes in viewing distance. Naturalness sensitivity exhibits scale-invariance for very large sizes with unfiltered textures (Figure 7), and over a wider range of sizes for low-pass filtered textures (Figure 8). That is, shifting task-relevant features through different retinal spatial frequencies does not degrade the ability of observers to use those features, as long as the features lie below the texture acuity limit. This scale-invariance is a common feature of suprathreshold form vision. Full or partial scale-invariance has been observed for many distinct form vision tasks, including texture discrimination (Dakin & Mareschal, 2000; Jamar & Koenderink, 1983; Joseph et al., 1997; Kingdom et al., 1995; Kingdom & Keeble, 1999; Sutter et al., 1995; van Meeteren & Barlow, 1981), texture segregation (Landy & Bergen, 1991; Nothdurft, 1985), contrast matching (Georgeson & Sullivan, 1975), symmetry detection (Dakin & Herbert, 1998; Rainville & Kingdom, 2002), spatial interval discrimination (Levi & Klein, 1990), reading (Legge et al., 1985), and object identification (Majaj et al., 2002; Oruç & Barton, 2010). Scale-invariance is also a feature of neural representations of form: cortical areas downstream of primary visual cortex encode object identity in a manner that is robust to changes in scale (Hong et al., 2016; Rust & DiCarlo, 2010). The scale-invariance of naturalness sensitivity supports the idea that, once early acuity limits are taken into account, visual form perception is highly invariant to changes in viewing distance.

### Cumulative sensitivity functions

Because of scale-invariance, naturalness sensitivity is more parsimoniously analyzed in terms of object spatial frequencies than retinal spatial frequencies. The cumulative sensitivity functions (Figure 13) show that, for all texture families studied, naturalness sensitivity rapidly improves with access to increasingly high object spatial frequencies. The curves also show differences between different texture families. Differences in the height of the curves illustrates that it is easier to distinguish naturalistic structure in some texture families than others (Freeman et al., 2013). Textures 1, 3, and 4 exhibit a qualitatively different shape (a rapid, saturating increase with spatial frequency) than textures 2 and 5 (a shallower slope with no saturation). We do not have a quantitative explanation for this difference, although inspection of the images (Figure 1E) suggests that the most salient naturalistic features in textures 1, 2, and 4 are “line-like,” while the most salient features in textures 2 and 5 are “blob-like.” Further study of these family-specific differences may help narrow down how specific image features are processed by mid-level visual mechanisms.

The cumulative sensitivity functions demonstrate that high object frequencies are *necessary* to achieve high levels of naturalness sensitivity. However, because these high frequency features were always presented in conjunction with lower frequency features, these experiments cannot demonstrate whether high frequency features are *sufficient* for high sensitivity. The model observer analysis shows that features in high spatial frequencies carry enough information to produce high sensitivity, but we do not know whether human sensitivity relies on the same underlying mechanisms as the model. Differences in sensitivity across texture families are most strongly predicted by “cross-scale” texture statistics that measure the co-occurrence of features in high and low object spatial frequency bands (see Appendix and Freeman et al., (2013)). This empirical result suggests that interactions between frequency bands may be a critical factor in driving high sensitivity. A definitive answer to this question will require measurements of sensitivity using bandpass filtered texture images that limit task-relevant information to a single spatial frequency band.

### Comparing human and model observers

We find that the naturalness sensitivity of human observers, on average, doubles for each additional octave-wide object spatial frequency band added to the image. We also find that a model observer trained on the same texture image set shows a doubling in sensitivity with spatial frequency. This particular rate of improvement may be related to the information-carrying capacity of each spatial frequency band. Parish & Sperling (1991) observed a similar rate of improvement with spatial frequency in the recognition of bandpass-filtered letters. They account for this rise by the increase in the number of effective samples in increasingly high spatial frequency bands: a factor of 4 for every additional octave-wide band. In an image of a given size, high-frequency features can independently covary at shorter distances, and thus at more locations, than low-frequency features. Each of these locations can vary independently in its local contrast and can (to a first approximation) be thought of as an independent sample. So long as these variations in local contrast range far above the contrast detection threshold, statistics measured in high-frequency bands will be averaged over more samples than statistics in low-frequency bands. This resulting reduction in sampling error predicts that sensitivity should be proportional to the square root of the increase in the number of independent samples. That is, each addition of an octave-wide frequency band should lead to a doubling in sensitivity. This matches the rate of improvement of both the cumulative sensitivity functions and of the model observer.

The observer model makes a number of strong predictions that can be empirically tested. For example, the model observer’s relatively high efficiency implies that much of the variance in psychophysical performance is intrinsically a property of the image set, rather than a limitation of the observer’s visual system. Because the model computes the strength of the naturalness signal for individual images, this hypothesis can directly tested by a “double-pass” experiment (Chin & Burge, 2020; van Meeteren & Barlow, 1981) that presents the same sets of images to the same observer multiple times. If observers respond with the same judgement to the same set of images across multiple repetitions, this would act as evidence that variance between the images was the true limiting factor on psychophysical performance.

The model observer also makes predictions about how strongly different texture statistics are weighted by the observer. By generating new texture images that parametrically vary in the relative strength of these different statistics, we can use the model observer as a fair baseline to judge which statistics most closely covary with human perception. In a similar vein, the current version of the model observer assumes that all locations within the aperture are weighted equally in the eventual decision. Psychophysical experiments using images that vary naturalness with spatial location could help confirm or refine this assumption. Using the model observer as a guide, these experiments could help constrain a more fully realized model of naturalness perception, and give insight into the mid-level processes that give rise to the perception of visual form.

### Connections to neurophysiology

The strong dependence of behavioral sensitivity on object spatial frequency raises the question of whether neural measurements of naturalness sensitivity exhibit a similar dependence on spatial frequency. Freeman et al. (2013) found no significant relationship between the preferred retinal spatial frequency of V2 neurons and the strength of their response to naturalistic structure. However, the model observer’s limitation by sampling error also predicts that having access to larger images (and thus more independent samples) should also drive higher sensitivity. This prediction is borne out by neural data: Larger images drive stronger naturalness modulation in V2 neurons than smaller images (Ziemba et al., 2018). The preferred spatial frequency of a visual neuron is typically inversely proportional to the size of its receptive field, such that neurons responsive to higher spatial frequencies tend to respond to smaller visual areas. In the model observer, these two factors effectively cancel, and thus predict no strong relationship between preferred spatial frequency and naturalness modulation. Rather, the model observer suggests that neural naturalness sensitivity might be best predicted by the ratio of a neuron’s preferred spatial frequency to its spatial extent: the number of spatial frequency cycles/receptive field diameter.

Our model observer demonstrates that each additional high spatial frequency band adds four times as many channels of information as the previous band, and that human observers extract this information with similar efficiency in all frequency bands. Both the results of the model observer and the structure of the Portilla & Simoncelli texture synthesis method suggest that an efficient visual system would, in a given region of space, distribute four times as many detectors in a high-frequency band than in a frequency band one octave lower. However, this prediction is inconsistent with neurophysiological data. Neuronal recordings in primary visual cortex typically indicate that at a given eccentricity, cells preferring high spatial frequencies are about as common as those preferring low spatial frequencies (De Valois et al., 1982). This suggests that visual cortex inefficiently samples the retinal image.

### Mid-level visual mechanisms

We conclude that our measurements of naturalness sensitivity are best explained by two factors that are *not* related to mid-level visual mechanisms. The first factor, the texture acuity limit, likely has its origin in contrast detection, and is thus a property of early visual processing. The second factor, the cumulative sensitivity functions of both human and model observers, shows that the strong dependence of naturalness sensitivity on high object spatial frequencies is primarily a property of the information available in the naturalistic texture image set, rather than a property of mid-level processing. In fact, our primary conclusions about mid-level processing relate to its constancy, rather than variation, with spatial frequency. The scale-invariance of naturalness sensitivity demonstrates that mid-level mechanisms are equally effective when the same image features are analyzed in different *retinal* spatial frequencies. The task efficiency measurements demonstrate that mid-level mechanisms show comparable effectiveness across different *object* spatial frequencies of the multi-scale texture images. This constancy suggests that naturalistic textures are a class of stimuli that is well-suited to engage multiple scales of mid-level processing at the same time.

## Acknowledgements

Thanks to Norma Graham, Mike Landy, and Corey Ziemba for helpful comments on the manuscript. Thanks to Eero Simoncelli, Jonathan Victor, and Tim Oleskiew for thought-provoking discussions on the results. Thanks to Sullivan Bacardo, Rui Diaz-Pacheco and Kiley Gan for help testing, maintaining, and running the experimental setup.

## Appendix: Model observer methods and comparison to Freeman et al., 2013

Our experimental results suggested that naturalness sensitivity relies on task-relevant information in high object spatial frequencies. To quantify which image features set the limits of naturalness discrimination, we built a model observer that produce estimates of naturalness sensitivity based only on the statistical properties of the texture images. While this model is not provably an ideal observer, it is computationally tractable and we expect its performance to be similar to that of an ideal observer.

The model observer computes the same statistics as the Portilla & Simoncelli texture synthesis method, then finds the combination of those statistics that optimally discriminates naturalistic texture from spectrally-matched noise. By computing discriminability for images of intermediate naturalness, we produced a relationship between discriminability and naturalness that allowed us to interpolate the model observer’s naturalness sensitivity. To measure how sensitivity changes with the removal of high-frequency image features, we grouped statistics by frequency band and systematically limited their availability to the discrimination algorithm. Finally, we used the model observer to investigate how different texture statistics contribute to naturalness discrimination.

### Model observer methods

The model observer begins by computing measurements of the same texture statistics used in the Portilla & Simoncelli synthesis procedure, with one difference. The model observer computes statistics from circularly apertured texture images (like the images used in the perceptual experiments) and restricts its spatial averaging of statistics to locations within the aperture. In contrast, the Portilla & Simoncelli procedure computes statistics on rectangular fields of texture that are assumed to have periodic image boundaries. As a result of these differences, for any individual texture image, the statistics measured by the model observer are not identical to the statistics used to synthesize the image. Rather, the measured statistics vary between different texture image examples. These measurements from the apertured images are a more realistic approximation of the information available to human observers when the texture images are realized on a physical screen.

We measured texture statistics for many different naturalistic texture images and spectrally-matched noise images (960 total per condition), then applied linear discriminant analysis to find the weighted sum of statistics that optimally separates naturalistic texture from noise. Because we had many more statistics than sample images, we used regularization and cross-validation to avoid overfitting. The full set of statistics, measured over the full ensemble of images, constitutes a matrix S of images by statistics. We define two matrices, *S_t_* and *S_n_*, for the statistics of the naturalistic texture and noise image sets, respectively. For each image set, we then measure the full covariance matrix for all statistics across all images in their set. We define Σ as the sum of these two covariance matrices:

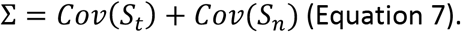

Linear discriminant analysis requires computing the inverse of this sum of covariance matrices. To regularize the inversion of the covariance matrix, we first define *Σ_diag_* as a matrix that is equal to Σ for all diagonal elements, and 0 otherwise. We then compute

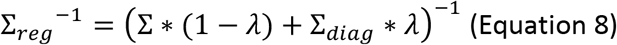

where *λ* is a regularization parameter bounded between 0 and 1. *λ*=0 computes the inverse covariance matrix with no regularization and *λ*=1 computes the inverse assuming that the statistics do not covary at all across images. The weights of the linear discriminant are then computed as

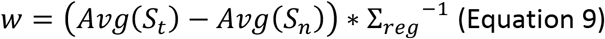

where *Avg*(*S_t_*) and *Avg*(*S_n_*) are the values of the statistics averaged over all images for the naturalistic texture and noise image sets respectively. To cross-validate our calculation of the weights, we split each image group into 5 partitions (192 images each), refitting our model each time holding out 1/5 of the data as a test set. We then applied the weights to the held out test data to compute a single, weighted sum discriminant value for each image. This resulted in two distributions of test set discriminant values: one for naturalistic images and one for noise images. We summarized the discriminability of these two distributions as a d-prime value

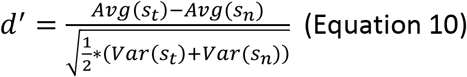

where *s_t_* is the distribution of the naturalistic discriminants and *s_n_* is the distribution of the noise discriminants. This analysis consistently found larger cross-validated d-prime values for *λ*>0 than for *λ*=0 (suggesting regularization helped avoid overfitting), and found stable d-prime values over a large range of *λ*s. For all analyses in this paper we set *λ* to 0.01, a value which achieved high levels of discriminability for all texture families. Our results did not qualitatively change for different choices of *λ*.

To compute the naturalness sensitivity of the model observer, we applied each texture family’s linear discriminant weights to statistics from images of intermediate naturalness. Specifically, we applied the training weights (found using fully naturalistic images) to test sets of statistics from images of intermediate naturalness. The training and test sets of images were segregated such that no seeds (the spectrally-matched noise images used to initialize the texture synthesis process) were shared between the two sets. Applying the weights to statistics from different naturalness levels allowed us to interpolate a relationship between discriminability (d-prime) and naturalness. Simulations of the match-to-sample task with Gaussian distributions reached threshold (75% correct) for d-prime values of 1.94. Accordingly, we defined the model observer’s threshold naturalness as the value corresponding to d-prime=1.94, and sensitivity as the inverse of threshold.

We repeated this process for “low-pass” observer models trained using subsets of the texture statistics limited to certain spatial frequency bands: only the lowest spatial frequency band, the 2 lowest, the 3 lowest, or the full set of all 4 frequency bands. For the purposes of all analyses we grouped cross-scale statistics into the higher of their two constituent frequency bands. These four frequency band subsets approximate the effect of low-pass filtering the texture images with corner object frequencies of 14, 28, 57, and 113 cycles/image, respectively. This process generated the model observer sensitivity measurements plotted in Figure 13.

We also repeated the process for “bandpass” observer models, each of which was limited to statistics from one of the four spatial frequency bands. As reported in the main text, bandpass model sensitivities were identical to low-pass model sensitivities when the single band of the bandpass model matched the highest band of the low-pass model. That is, the sensitivity of the highest frequency bandpass model was equivalent to that of the full (four band) low-pass model, the second highest bandpass was equivalent to the 3 lowest band model, etc. This calculation is complicated somewhat by the inclusion of cross-scale statistics which, contrary to the intention of the bandpass models, integrate information across different bands. We repeated these calculations for low-pass and bandpass models that did not have access to cross-scale statistics. Sensitivity was only slightly lower (average 8.8% decline) and was still equivalent between low-pass and bandpass conditions for all cases.

### Comparison of model observer to sensitivity regression from Freeman et. al. 2013

We used the model observer to determine how different subsets of statistics contributed to the discrimination of naturalistic texture from spectrally-matched noise. We trained multiple iterations of the model observer using either the full set of texture statistics (Figure 15A: “All”) or subsets of the statistics (e.g., cross-location statistics from simple cell-like linear filters, or cross-orientation statistics from complex cell-like energy filters, see Methods, image generation). We found energy covariance statistics resulted in stronger discrimination performance than linear covariance statistics (Figure 15A). We also found that cross-location statistics typically showed stronger discrimination performance than cross-orientation and cross-scale statistics.

**Figure 15.**
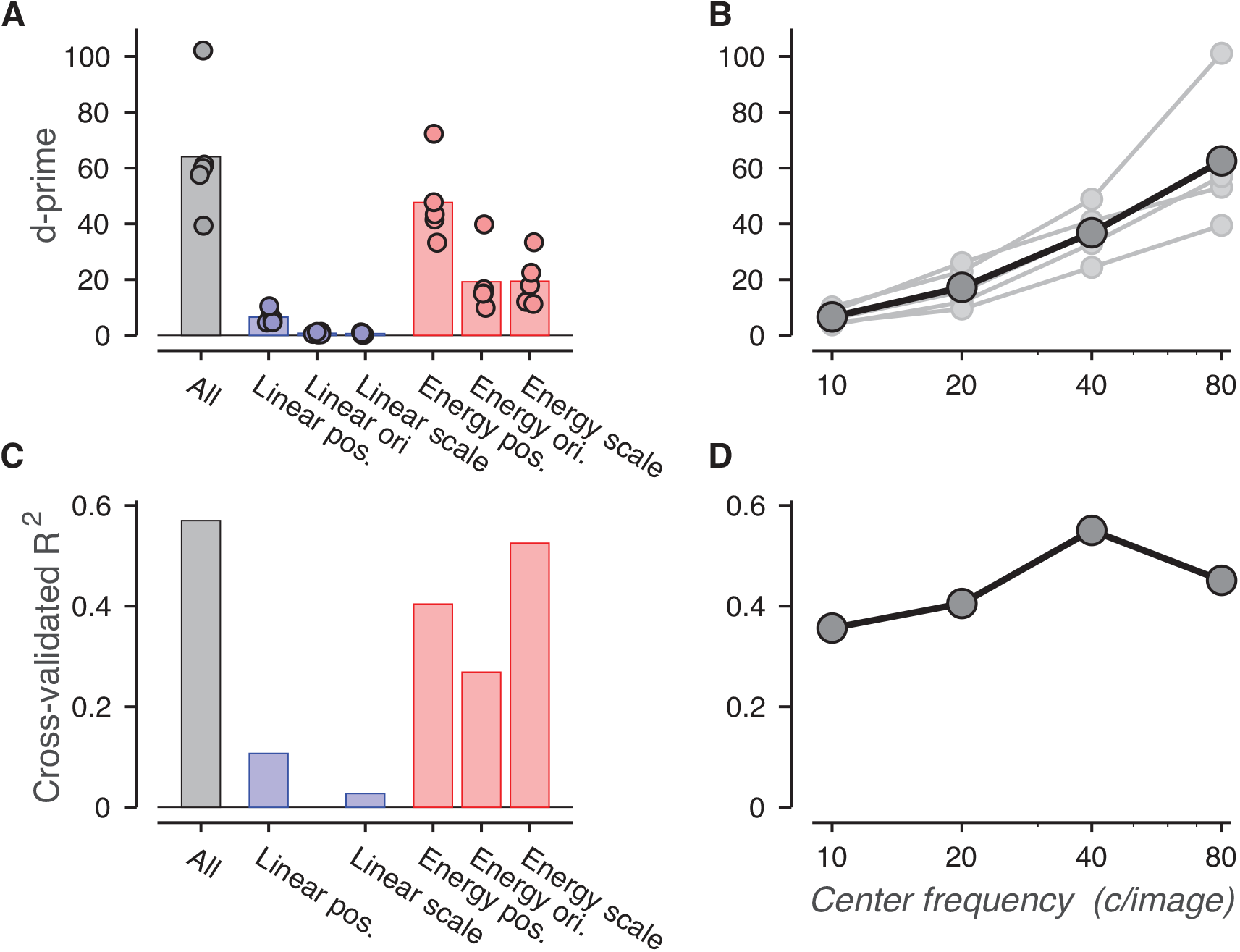
Naturalness sensitivity is best predicted by energy statistics and high object spatial frequency bands. A) Average model observer discriminability computed using different subsets of the texture statistics. Individual points denote the discriminability of individual texture families. B) Model observer discriminability computed using statistics from a single spatial frequency band, plotted versus object spatial frequency. The solid black line denotes the average over the five texture families. Each grey line denotes an individual texture family. C) R^2^ between naturalness sensitivity (444 texture families, data from Freeman et al. (2013)) and the predictions of a regression model. Each bar represents a regression model based on different subsets of the Portilla & Simoncelli texture statistics. D) R^2^ for predictions of human naturalness sensitivity computed using only Portilla & Simoncelli statistics that lie in a specific object spatial frequency band.

To examine the frequency-dependence of naturalness sensitivity, we plot discriminability from the bandpass model observers that only used subsets of statistics from one of the four different spatial frequency bands. Consistent with our psychophysical results, statistics in high frequency bands were much more effective at discriminating images than those in low frequency bands (Figure 15B).

We next compared the model observer’s discriminability measurements to the results of a parallel analysis from Freeman et al. (2013) that used multiple regression to determine which texture statistics most effectively predict human naturalness sensitivity. This analysis uses cross-validated multiple regression to determine how well different subsets of texture statistics predict naturalness sensitivity for 444 distinct texture families. We emphasize that this regression procedure is solving a qualitatively different task than that of the model observer. The model observer measures, for each individual texture family, the weighted sum of texture statistics that best discriminates naturalistic texture from noise. Each family uses a different set of weights and produces a different measurement of discriminability. In contrast, the regression analysis finds a single weighted sum of statistics that predicts differences in human naturalness sensitivity across many different texture families. Despite these different objectives, we suspected that both tasks would rely on the statistics that most strongly separate texture images from noise images.

Similarly to the model observer, the regression analysis predicted naturalness sensitivity more accurately using energy statistics than linear statistics (Figure 15C). However, unlike the model observer results, cross-scale energy statistics more accurately predicted naturalness sensitivity than cross-location and cross-orientation statistics (Figure 15C).

We believe this difference between the model observer and the regression analysis stems from differences between the two tasks. There are relatively few cross-scale statistics overall, and each of those statistics is typically a reliable marker of the overall strength of the naturalness signal in a texture family. Conversely, there are a much larger number of cross-location statistics, and their strength is more heterogeneous from texture family to family. A specific cross-location statistic that is stronger for naturalistic texture than noise in one family may have that relationship reversed in another family. As a result, the model observer may be learning family-specific patterns in the cross-location statistics that do not generalize to other texture families. To quantify this effect, we looked at how applying model observer weights trained on one texture family affected discriminability when those weights were applied to other families. We found that weights trained on cross-location or cross-orientation statistics were typically much less effective when those weights were applied to statistics from other texture families (average decline in d’: cross-location: 58%, cross-orientation: 64%). Conversely, weights trained on cross-scale statistics maintained high discriminability when applied to different families (average decline of 81%). Because the regression analysis relies on statistics that are strongly predictive across different families, it more heavily weights cross-scale statistics than cross-location statistics.

We next applied the regression analysis to statistics in different frequency bands. In contrast to the image observer results, we found that statistics in all four frequency bands were reasonably accurate at predicting perceptual sensitivity (Figure 15D). However, the predictive power of the second highest frequency band (40 c/image, R^2^=0.551) was essentially identical to that of the full model (R^2^=0.554), suggesting high levels of redundancy. Aside from the slight advantage of 40 c/image over other bands, we have trouble drawing strong conclusions from these data.

The small advantage of the 40 c/image statistics over the 80 c/image statistics contrasts with the model observer, which shows a significant advantage for the highest object spatial frequency band over all other spatial frequency bands. This discrepancy is likely due to the structure of the psychophysical task in Freeman et al. (2013). In this task, three texture images were presented simultaneously on the screen in the peripheral visual field. This task was run remotely on a large, crowd-sourced population of observers using Mechanical Turk, and as such the presentation size and distance could not be precisely controlled. Nonetheless, the target arrangement makes it likely that in most trials, the 80 c/image frequency band lay in retinal spatial frequency bands that were beyond the texture acuity limit of the observers. As such, it is reasonable that they are slightly less predictive than statistics in the highest *visible* spatial frequency band: 40 c/image.

## Notes

### Competing Interest Statement

The authors have declared no competing interest.

